# A porcine ex vivo lung perfusion model to investigate bacterial pathogenesis

**DOI:** 10.1101/708503

**Authors:** Amy Dumigan, Marianne Fitzgerald, Joana Sá Pessoa Graca Santos, Umar Hamid, Cecilia M. O’Kane, Danny F. McAuley, Jose A. Bengoechea

## Abstract

The use of animal infection models is essential to understand microbial pathogenesis and to develop and test treatments. Insects, and 2D and 3D tissue models are increasingly being used as surrogate for mammalian models. However, there are concerns whether these models recapitulate the complexity of host-pathogen interactions. Here, we developed the *ex vivo* lung perfusion (EVLP) model of infection using porcine lungs to investigate *Klebsiella pneumoniae*-triggered pneumonia as model of respiratory infections. The porcine EVLP model recapitulates features of *K. pneumoniae*-induced pneumonia lung injury. This model is also useful to assess the pathogenic potential of *K. pneumoniae* as we observed that the attenuated *Klebsiella* capsule mutant strain caused less pathological tissue damage with a concomitant decrease in the bacterial burden compare to lungs infected with the wild type. The porcine EVLP model allows assessment of inflammatory responses following infection; similar to the mouse pneumonia model, we observed an increase of *il-10* in the lungs infected with the wild type and an increase of *ifn-γ* in lungs infected with the capsule mutant. This model also allows monitoring phenotypes at the single-cell level. Wild-type *K. pneumoniae* skews macrophages towards an M2-like state. In vitro experiments probing pig bone marrow-derived macrophages uncovered the role of the M2 transcriptional factor STAT6, and that *Klebsiella*-induced *il10* expression is controlled by p38 and ERK. *Klebsiella*-induced macrophage polarization is dependent on the capsule. Altogether, this study support the utility of the EVLP model using pig lungs as platform to investigate the infection biology of respiratory pathogens.

**IMPORTANCE:** The implementation of infection models that approximate human disease is essential to understand infections and for testing new therapies before they enter into clinical stages. Rodents are used in most of pre-clinical studies, although the differences between mouse and man have fuelled the conclusion that murine studies are unreliable predictors of human outcomes. Here, we have developed a whole lung porcine model of infection using the established *ex vivo* lung perfusion (EVLP) system established to re-condition human lungs for transplant. As a proof-of-principle, we provide evidence demonstrating that infection of the porcine EVLP with the human pathogen *K. pneumoniae* recapitulates the known features of *Klebsiella*-triggered pneumonia. Moreover, our data revealed the porcine EVLP model is useful to reveal features of the virulence of *K. pneumoniae* including the manipulation of immune cells. Altogether, this study supports the utility of the EVLP model using pig lungs as surrogate host for assessing respiratory infections.

## INTRODUCTION

The use of animal infection models is essential to determine basic physiological principles, disease pathogenicity, identify virulence factors and to develop and test treatment strategies (1). The vast majority of immunology studies employ murine models, owing to availability of transgenic knockouts, reagents and established protocols. Therefore our current knowledge of the murine immune system far exceeds that of any other species. However, murine models have several limitations: there are significant differences between mice and humans in immune system development, activation, and response to challenge (2). Indeed mice share <10% genetic homology with human immune system (3). The increasing costs related to animal husbandry making large scale infection experiments expensive; and growing social concerns on the use of mice for biomedical experimentation despite the extensive and comprehensive animal welfare regulations in place, are additional drawbacks.

To circumvent these issues, alternative models of infection are being explored. Insects, including *Drosophila melanogaster* and *Galleria mellonella* (4), and the fish *Danio rerio* (5) are increasingly been used to investigate host-pathogen interactions. These models have proved successful in identifying virulence factors and to model features of the interaction between pathogens and the innate immune system. However, there are still concerns whether these infection models recapitulate the complex interactions between several immune cells, cytokines and chemokines and other soluble factors such as complement, and pathogens.

To address these issues, new infection models have been developed including 2D polarized epithelium, and 3D organoids of different tissues. These models still fall short of recapitulating the complex interactions between different cells as well as the structure of the organ. This study was initiated to establish a new infection model to investigate respiratory infections, the *ex vivo* lung perfusion (EVLP) model of infection using porcine lungs. Next to non-human primates, the domestic pig (*Sus scrofa domesticus*) has the closest genome and protein sequences compared to humans (6). Like humans, pigs are omnivores, share similar anatomy and physiology and have adaptive and innate immunes systems. Indeed, the porcine immune system is functionally more similar to the human immune system than that of mice, sharing >80% genetic homology (6). Notably, it is believed that experiments in pigs have more predictive therapeutic value than research carried out in rodents (7). The model developed herein facilitates the investigation of pathogen infection biology in a whole porcine lung receiving ventilation and perfusion in real time. This allows the investigation of the spatial distribution of infection, innate immune cell recruitment and activation, as well as histopathological changes. As a proof-of-concept, we have investigated whether this model recapitulates key features of *Klebsiella pneumoniae*-induced pneumonia.

*K. pneumoniae* is an important cause of nosocomial and community-acquired pneumonia. *Klebsiella* can readily spread between hospital patients with devastating results in immunocompromised individuals with mortality rates between 25-60% depending on the underlying condition (8). *K. pneumoniae* has been singled out by the World Health Organization as an urgent threat to human health due to the increasing isolation of multidrug resistant strains. A wealth of evidence obtained using the pneumonia mouse model demonstrates that clearance of *K. pneumoniae* relies on the activation of an inflammatory response which includes the activation of type I interferon (IFN)-controlled host defence responses (9, 10). Several studies have demonstrated the importance of alveolar macrophages and inflammatory monocytes in the containment and clearance of *K. pneumoniae* in the lungs (11–14). Conversely, this may suggest that a signature of *K. pneumoniae* infection biology is the attenuation of inflammatory responses and the subversion of macrophage-governed antimicrobial functions. Indeed, we and others have shown that, in sharp contrast to wild-type strains, attenuated mutant *Klebsiella* strains activate an inflammatory program, ultimately favouring their clearance (15–18). Furthermore, we have recently demonstrated that *K. pneumoniae* is able to survive intracellularly in mouse and human macrophages by preventing the fusion of lysosomes with the *Klebsiella* containing vacuoles (19).

Here, we report that the porcine EVLP infection model recapitulates key features of *Klebsiella*-triggered pneumonia. We present data showing that this model is also useful to assess the pathogenic potential of *K. pneumoniae* as we observed that the attenuated *Klebsiella* capsule mutant strain caused less pathological damage to the tissue with a concomitant decrease in the bacterial burden compare to lung infected with the wild-type strain. Finally, we present evidence demonstrating that *K. pneumoniae* skews macrophage polarization following infection in a STAT6 dependant manner.

## RESULTS

### *Ex vivo* lung porcine model of infection

Herein we have developed a whole lung porcine model of infection using the established EVLP model developed to re-condition human lungs that were marginal at meeting the lung retrieval criteria with the view to increase the lung donor pool for transplant (20). In this work, we have used one of the four commercially available clinical grade devices for EVLP, the Vivoline^®^ LS1 system. We selected a livestock porcine breed as they are readily available and been shown to best mimic animal variation reflective of human populations compared with wild breeds (7).

There are a number of essential details to consider when setting the porcine EVLP model. The quality of the organ is an essential factor, and researchers should carefully assess whether there are any macroscopic signs of damage/infection. The model uses 200 mL of autologous whole blood, which acts as a reservoir for immune cell recruitment and should be taken prior to lung retrieval. Lungs are removed from the pig and flushed with media through the pulmonary artery to remove blood. This is essential to avoid clotting. Lungs were then transferred to a sterile plastic bag on ice for transportation to the laboratory.

Unlike humans, pigs have an additional bronchus emerging from the trachea supplying the cranial lobe of the right lung (21). Therefore only left lungs were used in this investigation as they are immediately suitable for use on the LS1 system. However, preliminary experimentation revealed that by occluding the second bronchus on the right lung with a purse string suture, right lungs can also be used. A cannula is placed in the pulmonary artery and secured with surgical suture. An LS1 endobronchial tube is placed in the main bronchus and also secured with suture. To avoid inducing tissue damage and abnormal inflammatory responses the lungs should be warmed before any other manipulation, and the pulmonary perfusate and ventilation carefully managed in a gradual way. The lung is then connected to the LS1 system with perfusion but no ventilation and allowed to warm, ensure shunt is open at this time, once lung has reached 37°C, lung is inflated by hand using a bag-valve positive pressure ventilator assist device (Ambu-bag). The lung is then connected to a ventilator and receives 10 cm of continuous positive airway pressure (CPAP) with a mix of 95% oxygen with 5% CO_2_ (Figure 1A). A detailed description of the preparation of the lungs and set-up of the EVLP model is provided in the *Material and Methods* section. A schematic of a typical experimental design can be seen in Figure 1B.

**Figure 1:**
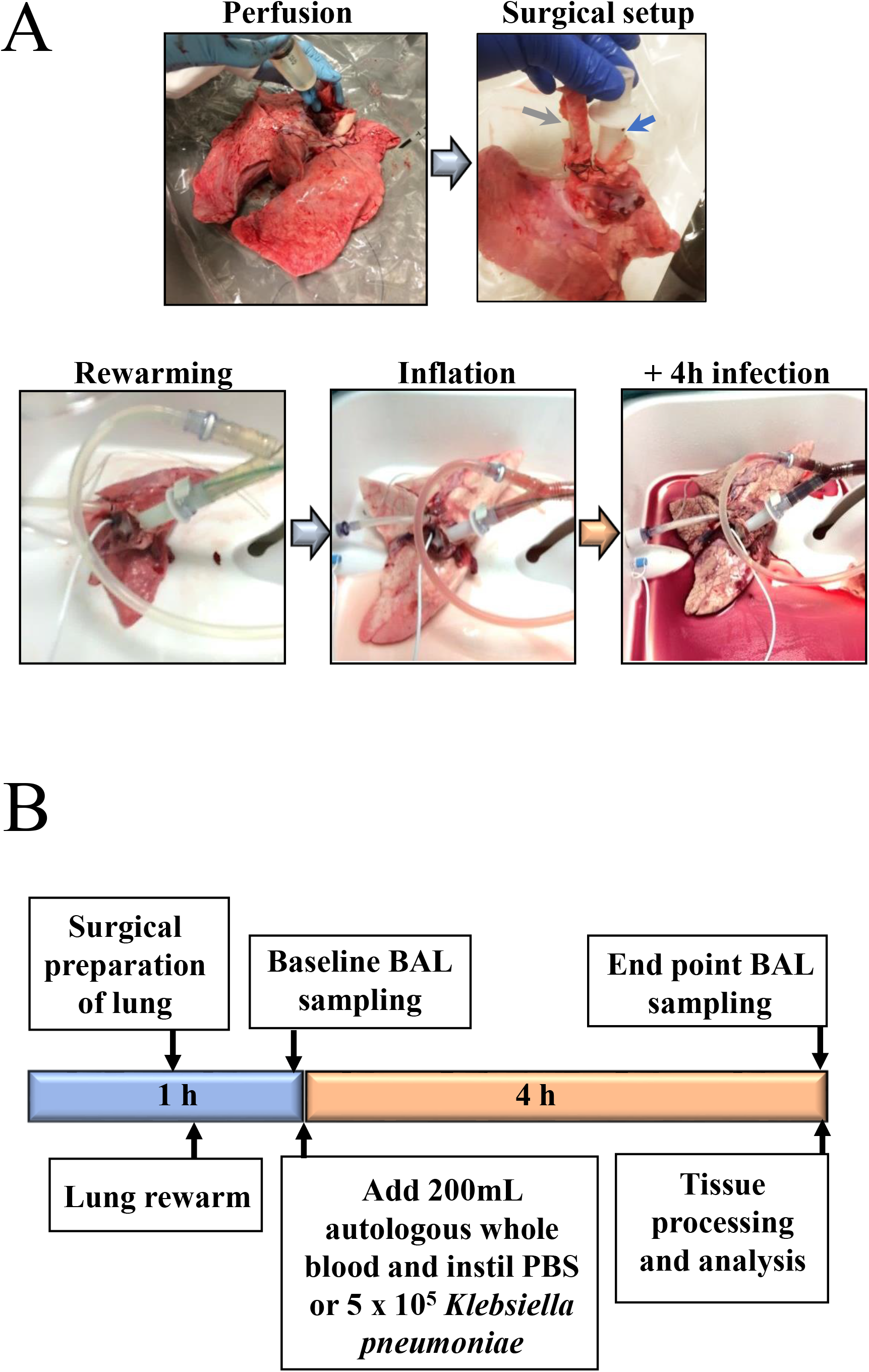
The porcine EVLP infection model. A. Images of lung during experimental process. Green arrow indicating LS1 ET tube in main bronchus and blue arrow indicating a catheter placed in pulmonary artery of left lung. B. Schematic describing the experimental design.

Preliminary experiments were carried to optimise the inoculum of *K. pneumoniae* 52.145 (hereafter Kp52145) and the time of infection based on macroscopic changes to the lungs. This *K. pneumoniae* strain clusters with those strains frequently associated with human infection and encodes all virulence functions significantly associated with invasive community-acquired disease in humans (22, 23). The virulence of this strain has been tested in several infection models including mice, rats, *G. mellonella*, and *Dyctiostlium discoideium* (24–27). An inoculum of 5 × 10^5^ CFU and 4 h infection period was selected in this study based on assessing macroscopic damage of lungs during infection, Once lung had been warmed to 37 ^o^C, a catheter was inserted into the caudal lobe of the lung and a baseline bronchoalveolar lavage (BAL) carried out. With catheter still in place, lungs received 5 mL of sterile PBS or inoculated with the bacterial inoculum. After 4 h of infection, a second BAL sample was collected and assessed for immune cell recruitment and protein levels. At the experimental endpoint, tissue samples were collected from the cranial, middle and caudal areas of the lung (Figure 1C) and analysed for oedema, bacterial colony forming units (CFU) and histology. Single samples were taken from the caudal lobe to assess immune cell recruitment (using flow cytometry) and gene transcription via real-time-qPCR (RT-qPCR).

### Tissue damage in the porcine EVLP model reflects hallmarks of *Klebsiella*-induced human pathology

Infection of porcine lungs with Kp52145 led to macroscopic damage after 4 h in stark contrast to PBS mock-infected lungs (Figure 2A). *K. penumoniae* capsule polysaccharide (CPS) is a well-characterized virulence factor of *Klebsiella* (28, 29)*. cps* mutant strains are avirulent in mammalian and non-mammalian models of disease (24, 25, 28, 29). To determine the sensitivity of the porcine model, lungs were infected with 5 × 10^5^ CFU in 5 mL PBS of strain Kp52145-Δ*wca*_*K2*_ As shown in Figure 2A, infection with the *cps* mutant resulted in limited macroscopic damage, suggesting that the *cps* also plays a crucial role in infection biology of *K. pneumoniae* in the porcine EVLP infection model.

**Figure 2:**
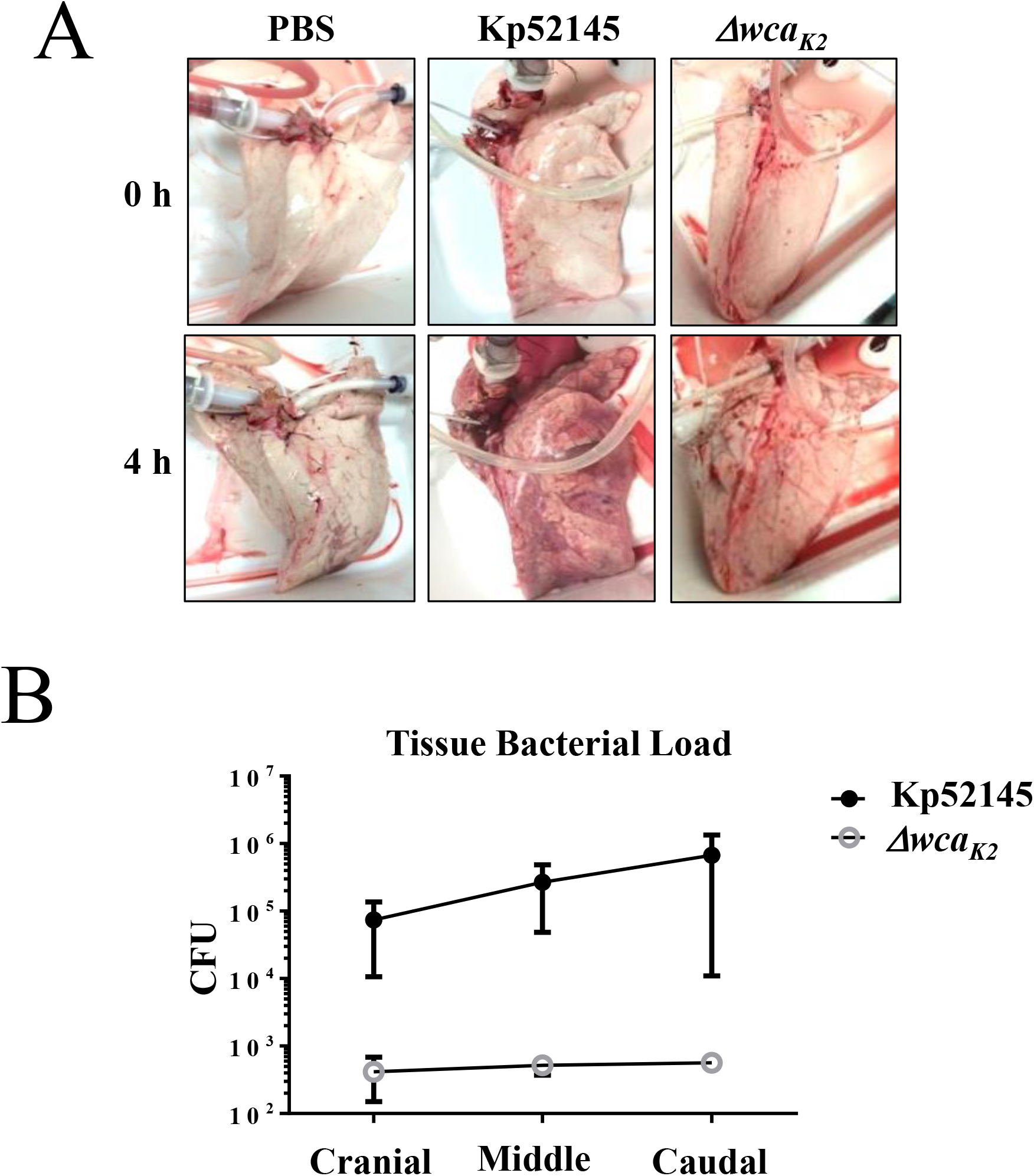
Infection of whole lungs with *K. pneumoniae* induces lung damage. A. Images of macroscopic damage of the lungs before and after infection with *K. pneumoniae* 52.145 (Kp52145) and the isogenic *cps* mutant, strain 52145-Δ*wca*_*K2*_. B. Bacterial load (CFU per gr of tissue) across different sections of the lungs infected with *K. pneumoniae* 52.145 (Kp52145) and the isogenic *cps* mutant, strain 52145-Δ*wca*_*K2*_. Values are presented as the mean ± SEM of three independent experiments.

To establish whether the macroscopic damage in the lungs infected with the wild-type strain was associated with higher bacterial burden in the tissue, samples were collected across the lung, as shown in Figure 1C, homogenized and the number of CFUs per gram of tissue determined. Indeed, the bacterial burden was three logs higher in lungs infected with the wild-type strain than in those infected with the *cps* mutant. Interestingly, despite inoculum being introduced in the caudal lobe, bacterial burden was homogenously distributed across the lung (Figure 2B).

Histological analysis of porcine tissues was carried out based on parameters of acute respiratory distress syndrome (ARDS) in animal models as defined by the American Thoracic Society. Pathogenic hallmarks of lung injury include: thickening of alveolar septa and infiltration of proteinaceous debris, red blood cells (haemorrhage) and immune cells including neutrophils into the alveolar space (neutrophilic alveolitis) (30). Analysis of lung stained sections with hematoxilin-eosine revealed signs of injury in lungs infected, although injury was more severe in those lungs infected with Kp52145 (Figure 3A). This was further confirmed by analysis of alveolar septal thickness (Figure 3B). This measurement revealed significant thickening of alveolar septal membranes in lungs infected with Kp52145. Interestingly, infection with the *cps* mutant strain induced significantly enhanced alveolar septal thickening compared to PBS controls, however this damage was significantly reduced when compared to Kp52145-infected lungs (Figure 3B). One hallmark of *K. pneumoniae*-triggered necrotising pneumonia is the presence of cherry red (blood streaked) sputum, i.e. haemorrhage. Haemorrhage is clearly evident both macroscopically (Figure 2A) and microscopically (Figure 3A) in lungs infected with Kp52145, and significantly reduced in the lungs infected with the *cps* mutant. The presence of intra-alveolar haemorrhage was assigned a score of 0, 1, 2 or 3 based on a semi-quantitative assessment of none, mild, moderate or severe. Scoring confirmed significantly enhanced haemorrhage in lungs infected with Kp52145 compared to the lungs PBS-mock infected and infected with the *cps* mutant (Figure 3C). Haemorrhage was accompanied by presence of inflammatory immune cells within the alveolar space. The number of nucleated cells in the alveolar space was quantified, and it was significantly higher in the lungs infected with Kp52145 than in those infected with the *cps* mutant or PBS-mock infected (Figure 3D). The presence of proteinaceous debris was significantly higher in the infected lungs compare to those PBS-mock infected. However, proteinaceous debris was significantly higher in lungs infected with Kp52145 than in those infected with the *cps* mutant (Figure 3E).

**Figure 3:**
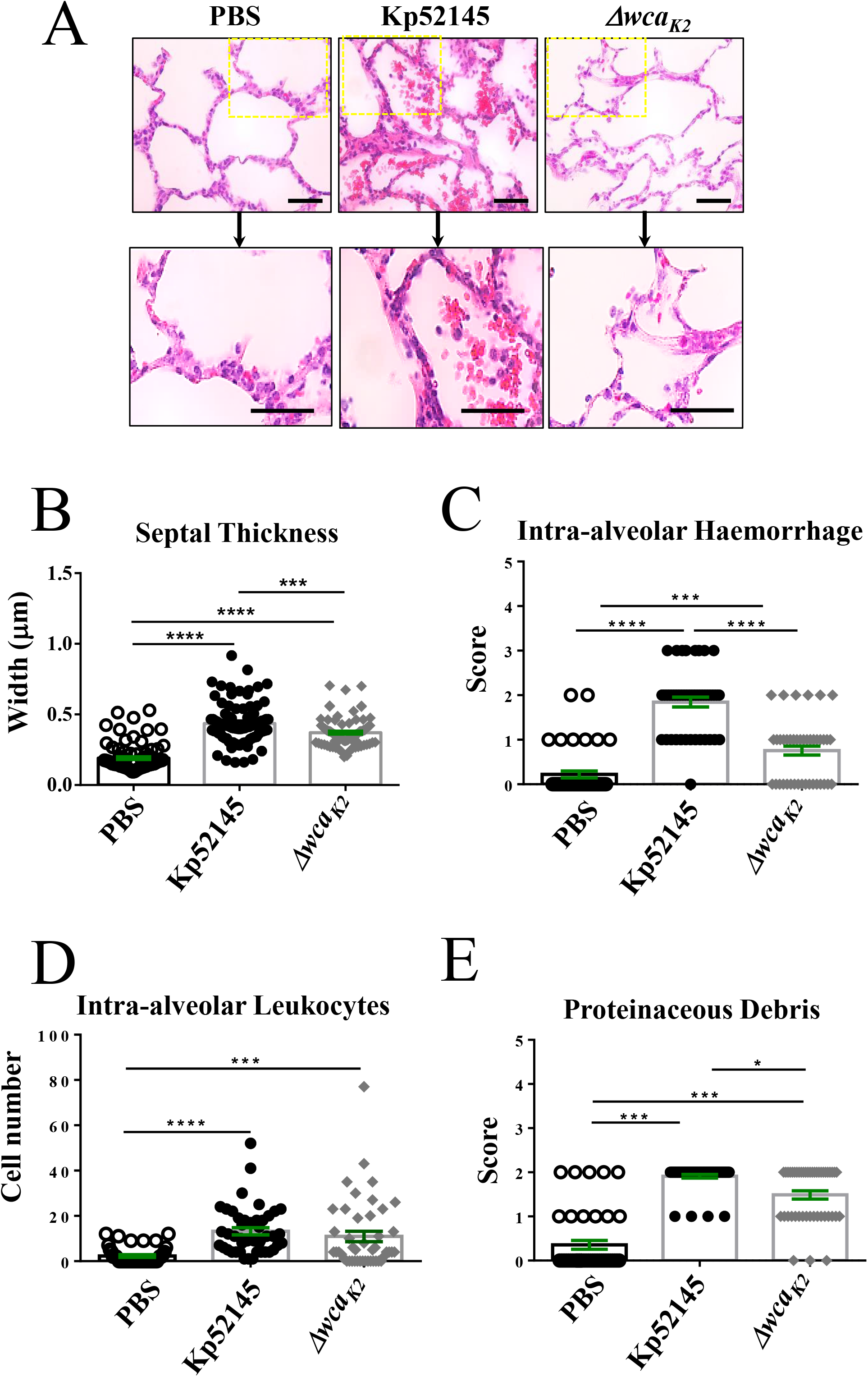
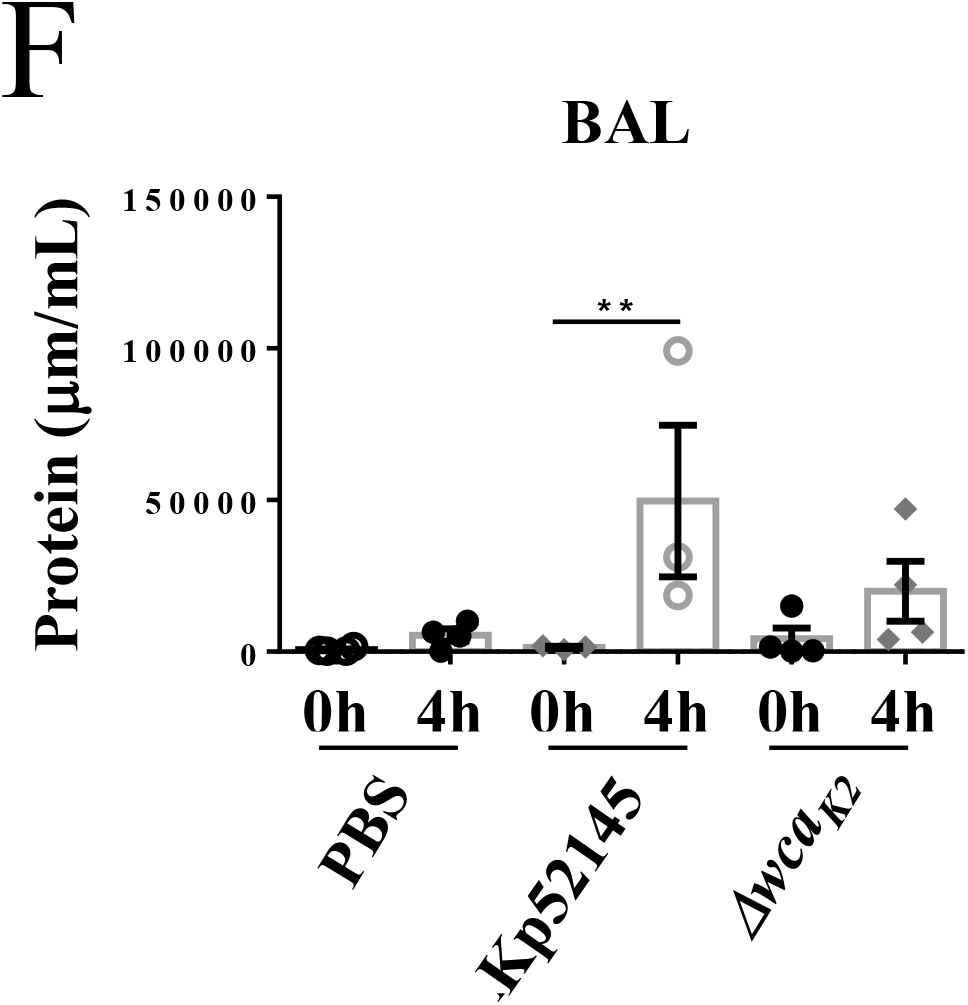
Porcine EVLP model recapitulates clinical hallmarks of *K. pneumoniae* induced pneumonia. A. Haematoxylin and eosin staining of porcine lung samples (x 400 magnification) from lungs mock-infected (PBS), and infected with *K. pneumoniae* 52.145 (Kp52145) and the isogenic *cps* mutant, strain 52145-Δ*wca*_*K2*_. B. Alveolar septal thickness was measured using ImageJ software. Each dot represents an average of three alveolar thicknesses per image, corresponding to three sections per lung across three experimental replicates from lungs mock-infected (PBS), and infected with *K. pneumoniae* 52.145 (Kp52145) and the isogenic *cps* mutant, strain 52145-Δ*wca*_*K2*_.(Δ*wca*_*K2*_). C. Intra-alveolar haemorrhage was scored per image whereby 0, 1,2, and 3 represent none, mild, moderate and severe levels of red blood corpuscles within the alveolar space from lungs mock-infected (PBS), and infected with *K. pneumoniae* 52.145 (Kp52145) and the isogenic *cps* mutant, strain 52145-Δ*wca*_*K2*_.(Δ*wca*_*K2*_). D. Number of nucleated cells evident in the alveolar space per image from lungs mock-infected (PBS), and infected with *K. pneumoniae* 52.145 (Kp52145) and the isogenic *cps* mutant, strain 52145-Δ*wca*_*K2*_.(Δ*wca*_*K2*_). E. Scoring of proteinaceous debris in the alveolar space from lungs mock-infected (PBS), and infected with *K. pneumoniae* 52.145 (Kp52145) and the isogenic *cps* mutant, strain 52145-Δ*wca*_*K2*_.(Δ*wca*_*K2*_). F. Protein levels at baseline and endpoint BAL samples from whole lungs mock-infected (PBS), and infected with *K. pneumoniae* 52.145 (Kp52145) and the isogenic *cps* mutant, strain 52145-Δ*wca*_*K2*_.(Δ*wca*_*K2*_). Statistical analysis was carried out using one-way ANOVA, ****p<0.0001, ***P < 0.001; **P < 0.01. Error bars are standard error of mean.

Further supporting that infection with Kp52145 was associated with an increase in lung injury, thirty five-fold increase in the total levels of BAL protein was found in the lungs infected with the wild-type strain. There were no differences in the total BAL protein between lungs infected with the *cps* mutant and PBS-mock infected (Figure 3F). These findings suggest that infection with the wild-type strain affected alveolar epithelial-endothelial barrier function.

Collectively, these findings demonstrate that the porcine EVLP model recapitulates features of *K. pneumoniae*-induced pneumonia lung injury. Furthermore, our results demonstrate that this model is useful to assess the virulence of *K. pneumoniae* since the *cps* mutant, known to be attenuated in other infection models (24, 25, 28, 29), was also attenuated in the porcine EVLP model.

### Innate Immune cell recruitment in *K. pneumoniae* EVLP model

We next sought to investigate innate immune response to *K. pneumoniae* infection in the porcine EVLP model. 100 μg of tissue were removed from caudal lobe and homogenised. Red blood cells were removed from BAL and tissue samples using ammonium-chloride-potassium lysis buffer. Samples were then stained for innate immune cells using purified anti-pig antibodies conjugated with fluorophores and analysed by flow cytometry. CD11R3 has a similar expression pattern to the human CD11b marker, being expressed on pig monocytes and alveolar macrophages, but not on lymphocytes, eythrocytes or platelets (31, 32) and was used to assess macrophages. Porcine CD172a, a marker of dendritic cells (33), and a porcine specific granulocyte marker clone 6D10 (Bio-Rad) to identify neutrophils were also used (31, 32). Lungs infected with Kp52145 showed an increase in the number of macrophages in tissue (Figure 4A). Macrophages are presented as percentage single CD11R3+ cells (gating strategy and representative dot plots supplied in Supplementary Figure 2A). Neutrophils were identified using a porcine granulocyte marker (clone 6D10) in a similar fashion (Supplementary Figure 2B) and were shown to be increased in density in 4 h BAL in Kp52145 infected experimental group (Figure 4B). No significant change was observed in CD11R3-CD172+ dendritic cells (Figure 4C) (gating strategy described in Supplementary Figure 2C). The number of macrophages and neutrophils in the tissue and BAL from lungs infected with the *cps* mutant were lower than those found in the wild-type-infected lungs and closer to the number found in PBS-mock infected lungs (Figure 4B). Enhanced macrophage and neutrophil recruitment in Kp52145 infected BAL samples and lung tissue respectively correlates with injury observed in histological analysis (Figure 3A-EF).

**Figure 4:**
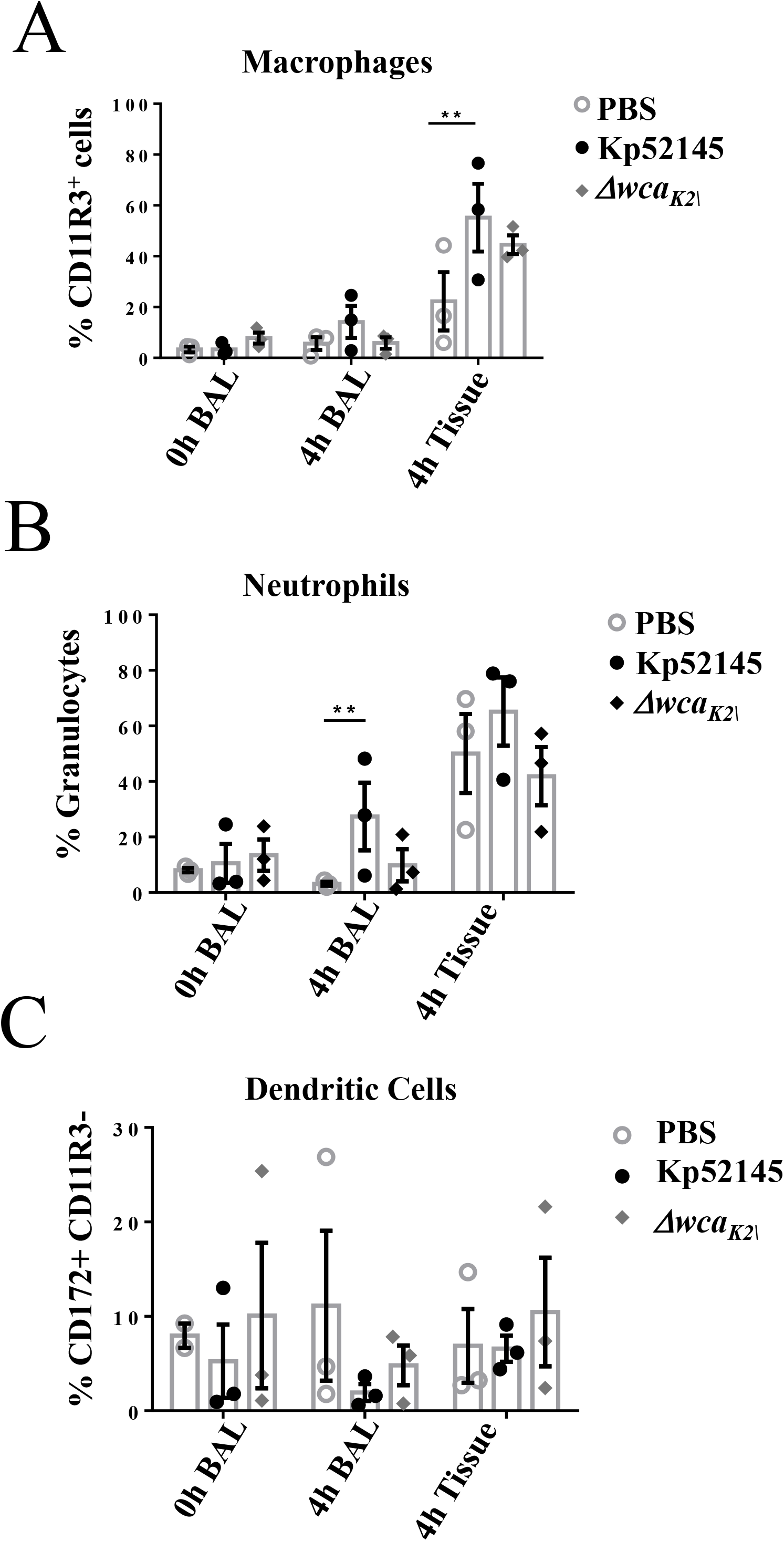
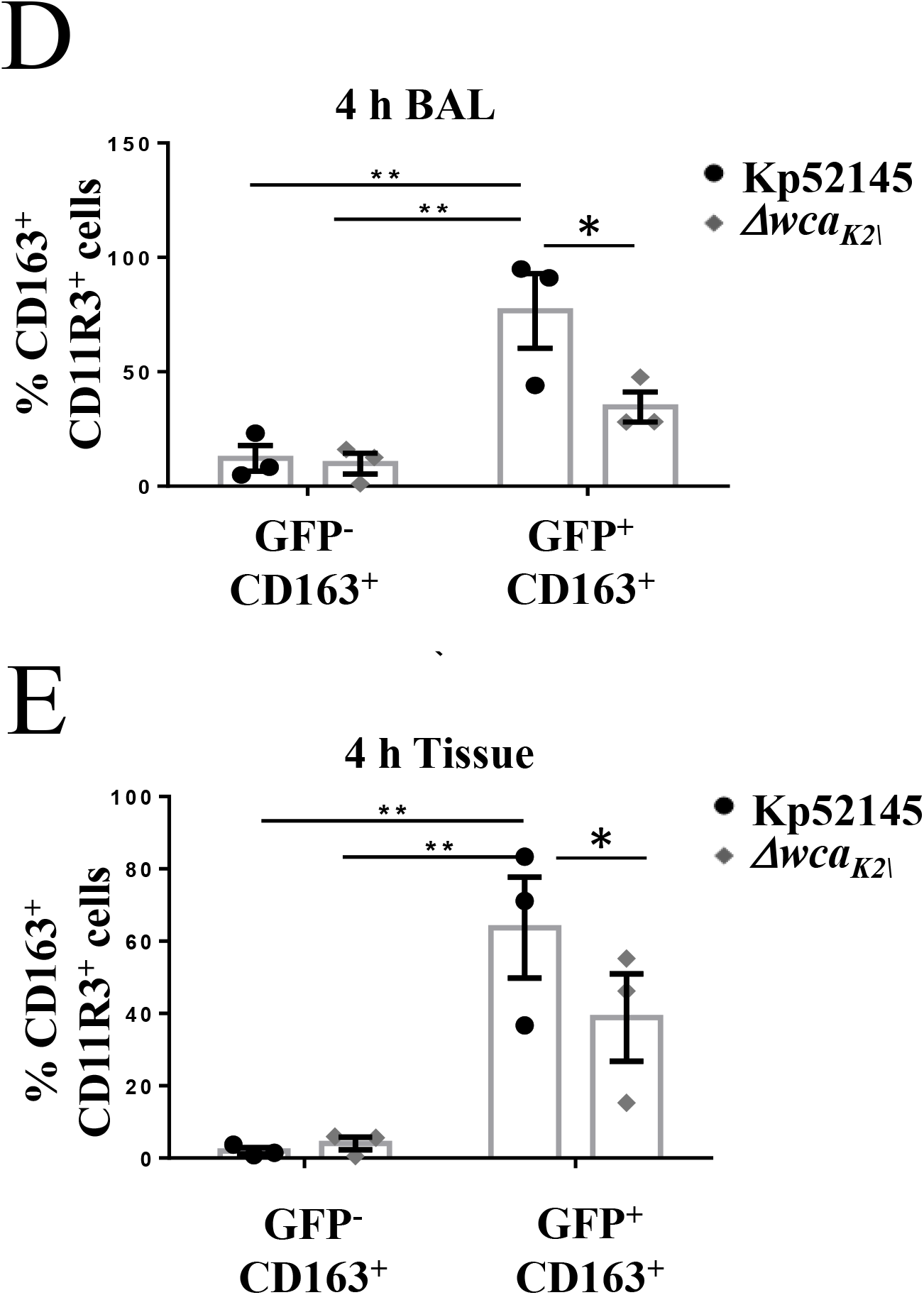
Innate cells recruitment in *K. pneumoniae*-infected porcine EVLP model. A. % CD11R3+ macrophages in baseline (0 h) and endpoint (4 h post treatment) in BAL samples and tissue from caudal lobe of mock-infected (PBS), and infected with *K. pneumoniae* 52.145 (Kp52145) and the isogenic *cps* mutant, strain 52145-Δ*wca*_*K2*_. B. % Granulocytes in baseline (0 h) and endpoint (4 h post treatment) in BAL samples and tissue from caudal lobe of mock-infected (PBS), and infected with *K. pneumoniae* 52.145 (Kp52145) and the isogenic *cps* mutant, strain 52145-Δ*wca*_*K2*_. C. % CD172+ dendritic cells in baseline (0 h) and endpoint (4 h post treatment) in BAL samples and tissue from caudal lobe of mock-infected (PBS), and infected with *K. pneumoniae* 52.145 (Kp52145) and the isogenic *cps* mutant, strain 52145-Δ*wca*_*K2*_. Percentage of CD11R3^+^ macrophages positive for CD163 expression associated (GFP^+^) or not (GFP^-^) with *K. pneumoniae* 52.145 (black dots)) and the isogenic *cps* mutant, strain 52145-Δ*wca*_*K2*_ (white dots) harbouring plasmid pFPV25.1Cm in BAL (D) and tissue (E). In all panels, values are represented as standard error of mean of three independent experiments; **p < 0.001,*p < 0.05 determined by unpaired t test.

The presence of bacteria in tissues is associated with macrophage reprogramming (34). M1 (classical) polarization is associated with protection during acute infections, whereas M2 (alternative) programme is linked to the resolution of inflammation and tissue regeneration (34). Therefore, we sought to establish whether *K. pneumoniae* infection could be linked to a macrophage switch in polarization. To investigate this possibility, we assessed the levels of the known M2 macrophage marker CD163, an iron scavenger receptor, in infected macrophages (35). Infections were carried out with bacteria expressing GFP to assess CD163 levels in cells with and without associated bacteria. Flow cytometry experiments showed that the levels of CD163 were significantly higher in those macrophages associated with Kp52145 (CD11R3+GFP+CD163+) than in those without bacteria (CD11R3+GFP-CD163+) (Figure 4D) (gating strategy can be found in Supplementary Figure 2D). Interestingly, when infections were done with the *cps* mutant, the levels of CD163 were significantly lower in macrophages associated with the mutant than in those associated with the wild-type strain. (Figure 4D and E), suggesting that the CPS may contribute to expression of CD163 on macrophages in *K. pneumoniae*-infected lungs.

### *K. pneumoniae*-induced inflammation in the porcine EVLP model

To further investigate the host response to *K. pneumoniae* in the porcine EVLP model, we analysed the expression of several inflammation-associated cytokines and chemokines by RT-qPCR from samples collected from the caudal lobe of lungs. Higher levels of *il-6* and *il-12* were detected in the lungs infected with Kp52145 than in those infected with the *cps* mutant or PBS-mock infected (Figure 5A and B). In contrast, the levels of *il-8,* and *ifn-γ* were significantly higher in the lungs infected with the *cps* mutant than in those infected with Kp52145 (Figure 5C and D). Mice deficient in IFN-*γ* production suffer greater mortality from *K. pneumoniae* infection (36–39). The higher levels of IFN*γ* that are produced during *cps* mutant infection in the EVLP model are likely a result of the high rate of clearance of the capsule mutant strain.

**Figure 5:**
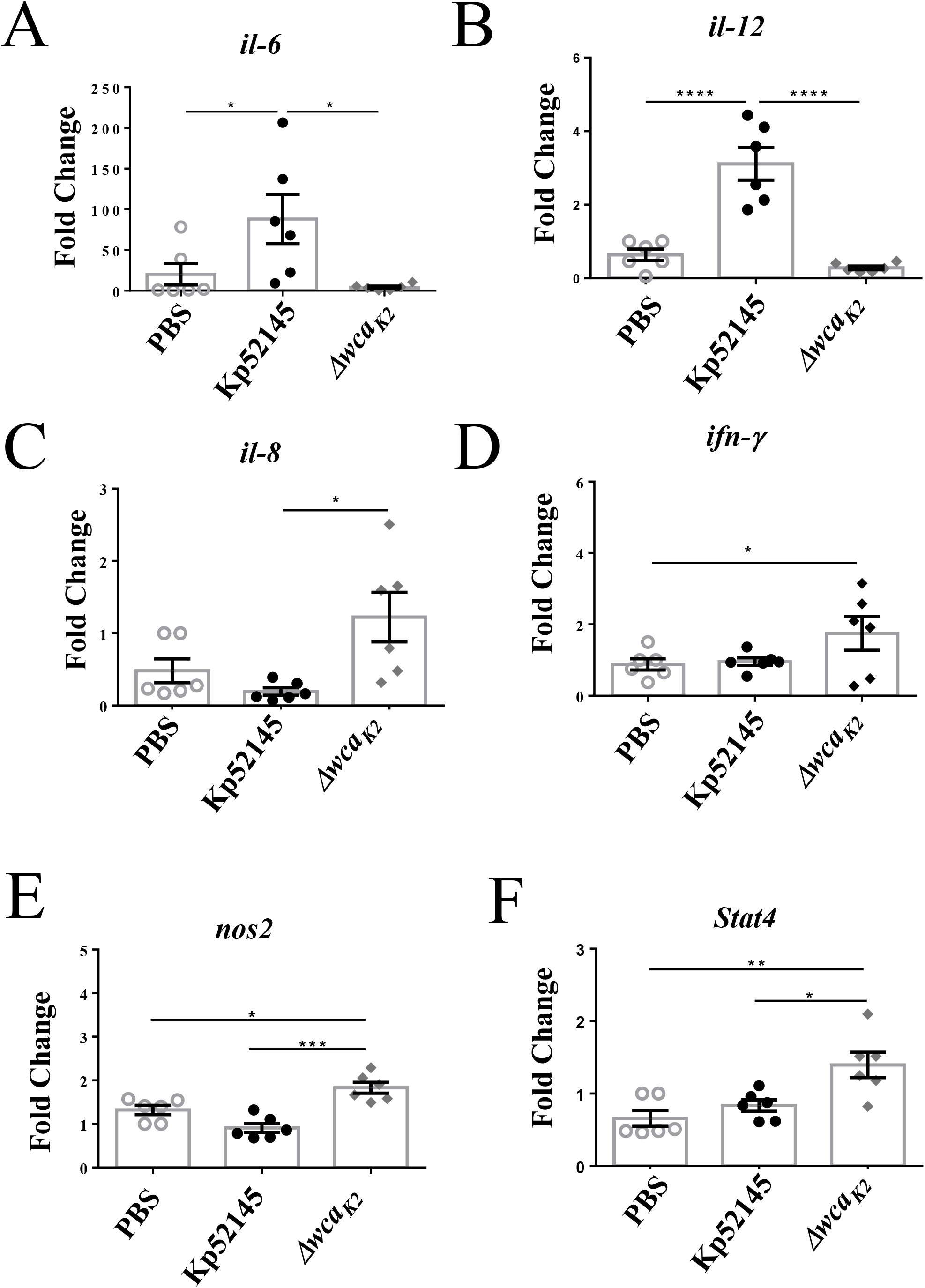
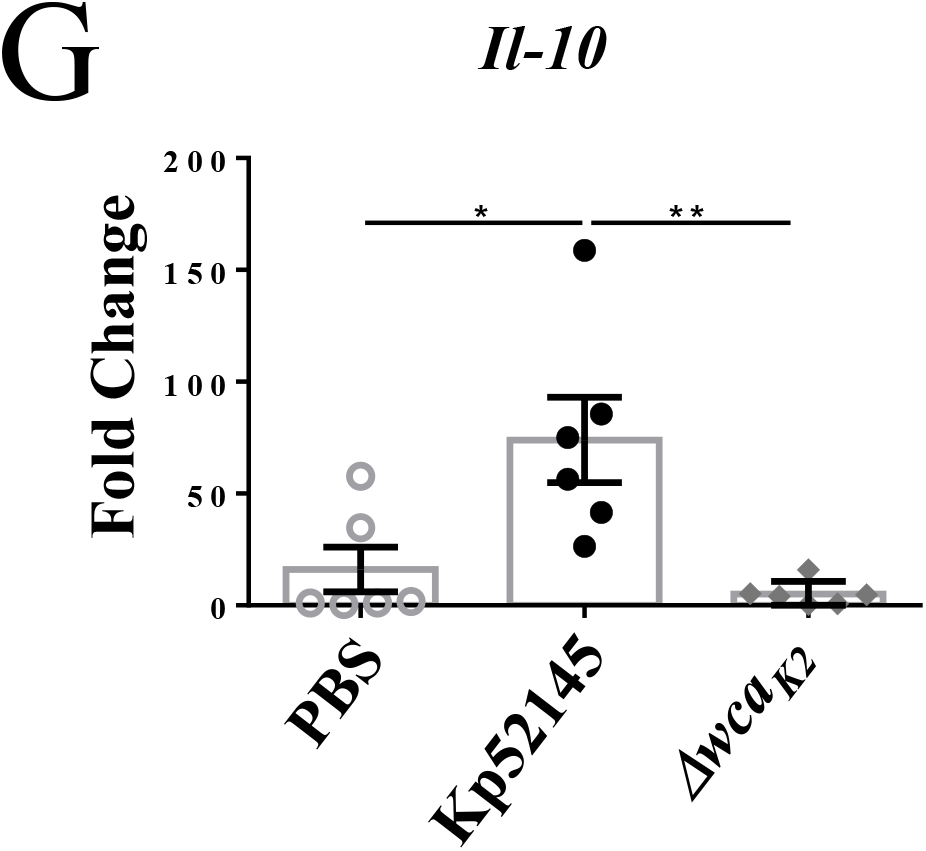
−*K. pneumoniae* induced inflammation in the porcine EVLP model. mRNA levels in lung tissues mock-infected (PBS), and infected with *K. pneumoniae* 52.145 (Kp52145) and the isogenic *cps* mutant, strain 52145-Δ*wca*_*K2*_.(Δ*wca*_*K2*_) assessed by RT-qPCR: A. interleukin (il)−6, B. *il-12*, C. *il-8,* D. *ifn-γ* E. *nos2*, F. *stat-4*, G. *il-10*. Values are presented as the mean ± SEM of three independent experiments measured in duplicate. ****p<0.0001, ***p<0.001, **p<0.01, *p<0.05 for the indicated comparisons using one-way ANOVA with Bonferroni correction.

The expression levels of *nos2* and *stat4* were also significantly higher in the lungs infected with the *cps* mutant than in those infected with the wild-type strain which were similar to those lungs PBS-mock infected (Figure 5 E and F). These markers have been associated with M1 polarized macrophages (35). We observed a significant increase in the levels of the anti-inflammatory cytokine *il-10* only in the lungs infected with Kp52145 (Figure 5G). Similar observation has been reported previously in the mouse pneumonia model (40, 41). Notably, enhanced production of *il-10* is one of the features of M2 polarized macrophages which are associated with resolution of inflammation (34, 35, 42).

Collectively, these findings demonstrate that the porcine EVLP model is useful to assess inflammatory responses following infection. By assessing *Klebsiella*-induced responses, our results infer that wild-type *K. pneumoniae* may modulate macrophage polarization towards M2 state.

### *K. pneumoniae* drives macrophage polarisation in a STAT6-dependent manner

To further investigate whether *K. pneumoniae* governs macrophage polarization, we established a method to generate porcine bone marrow-derived macrophages (pBMDMs). We next sought to determine whether *K. pneumoniae* skews the polarization of pBMDMs. Infection of pBMDMs with Kp52145 resulted in a significant upregulation of the surface expression of the M2 marker CD163 as detected by flow cytometry (Figure 6A). This finding is in perfect agreement with the results obtained infecting the porcine EVLP model. STAT6 is a well-established transcription factor regulating M2 macrophage polarization (43, 44). Therefore, we sought to determine whether *K. pneumoniae* activates STAT6 to govern macrophage polarization in pBMDMs. Immunoblotting experiments revealed that Kp52145 induced the phosphorylation of STAT6 in pBMDMs (Figure 6B). Phosphorylation of STAT6 is essential for its nuclear translation to control the transcription of STAT6-induced genes (45, 46).To establish whether *Klebsiella*-induced macrophage polarization is STAT6-dependent, we followed a pharmacologic approach probing the STAT6 inhibitor AS1517499 (47). Transcriptional analysis showed that *Klebsiella*-induced expression of the M2 markers, *cd163* and *arginase-1*was ablated in cells pre-treated with the STAT6 inhibitor (Figure 6C and D), demonstrating that *Klebsiella* induction of M2 markers is STAT6 dependent. Interestingly, and in agreement with our previous findings suggesting that the CPS could be required for *Klebsiella*-triggered macrophage polarization, the *cps* mutant did not induce the phosphorylation of STAT6 (Figure 6E). As we anticipated, the *cps* mutant did not induce the expression the M2 markers *arg-1* and *cd163* in pBMDMs (Figure 6F and G). Furthermore, the *cps* mutant induced the expression of the M1 markers *stat4* and *nos2* (Supplementary Figure 3).

Kp52145 also upregulated the transcription of the anti-inflammatory cytokine and M2 marker *il-10* in pBMDM (Figure 7A), indicating the *il-10* expression observed in porcine EVLP tissues infected with Kp52145 (Figure 5G) could be derived from macrophages. This increased expression was not dependent on STAT6 because the STAT6 inhibitor did not reduce the expression of *il-10* (Figure 7A). In mouse and human macrophages, the transcription of *il-10* is regulated by STAT3 (48). Immunoblotting analysis confirmed the activation of STAT3 in *Klebsiella*-infected pBMDMs (Figure 7B). MAP kinases p38 and ERK are known to control the expression of IL10 in mouse and human macrophages (48).

**Figure 6:**
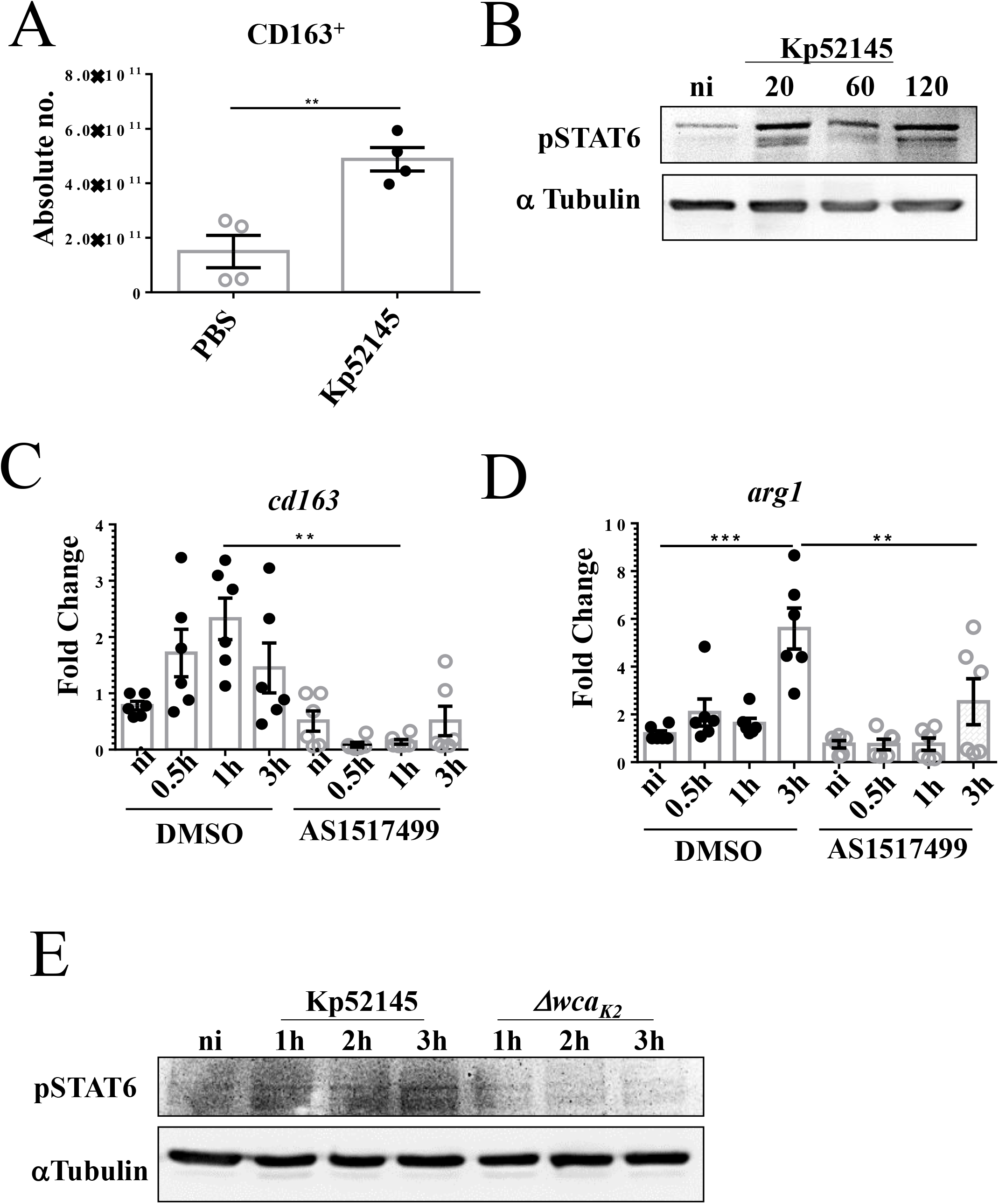
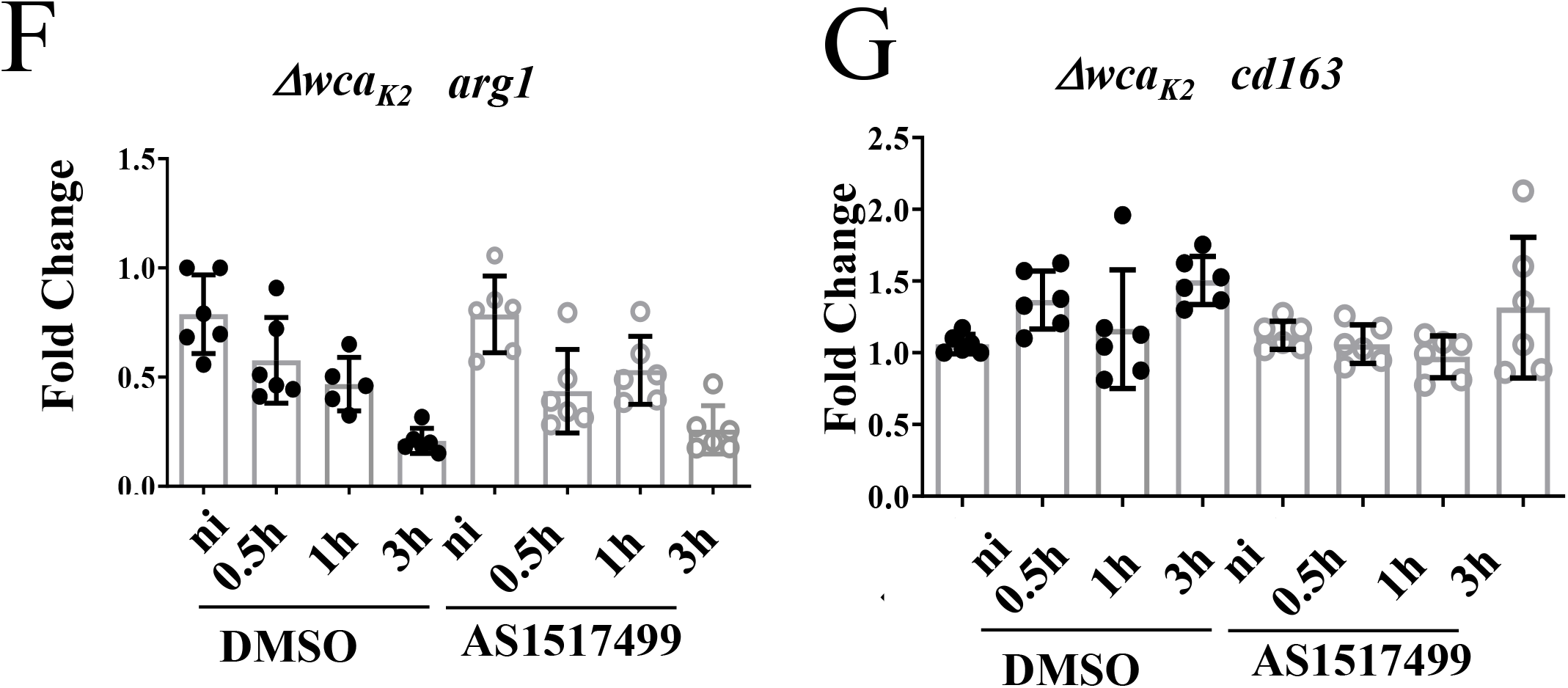
*K. pneumoniae* drives macrophage polarisation in a STAT6-dependent manner. A. CD163 surface expression in PBS or Kp52145-infected pBMDMs by flow cytometry. Values are shown as standard error of mean of two independent experiments in duplicate. **, p<0.01 determined by unpaired Student’s -test. B. Immunoblotting analysis of phosphorylation of STAT6 (PSTAT6) and tubulin in lysates of pBMDMs infected with Kp52145 for the indicated times or left uninfected (ni). Data is representative of three independent experiments. C. *cd163* levels in pBMDMs non-infected (ni) or infected with Kp52145 pre-treated with STAT6 inhibitor (AS1517499, 50nM 2 h prior to infection) or DMSO vehicle control. Values are shown as standard error of mean of three independent experiments. D. Arginase-1 levels in pBMDMs non-infected (ni) or infected with Kp52145 pre-treated with STAT6 inhibitor (AS 1517499, 50nM 2 h prior to infection) or DMSO vehicle control. Values are shown as standard error of mean of three independent experiments. E. Immunoblotting analysis of phosphorylation of STAT6 (PSTAT6) and tubulin in lysates of pBMDMs infected with Kp52145 and the isogenic *cps* mutant, strain 52145-Δ*wca*_*K2*_ for the indicated times or left uninfected (ni). Data is representative of three independent experiments. F. *cd163* levels in pBMDMs non-infected (ni) or infected with the *cps* mutant, strain 52145-Δ*wca*_*K2*_, pre-treated with STAT6 inhibitor (AS1517499, 50nM 2 h prior to infection) or DMSO vehicle control. Values are shown as standard error of mean of three independent experiments in duplicate. G. *arginase-1* levels in pBMDMs non-infected (ni) or infected with *cps* mutant, strain 52145-Δ*wca*_*K2*_, pre-treated with STAT6 inhibitor (AS 1517499, 50nM 2 h prior to infection) or DMSO vehicle control. Values are shown as standard error of mean of three independent experiments in duplicate. In panels C, and D, ***p<0.001, **p<0.01, for the indicated comparisons using one-way ANOVA with Bonferroni correction.

**Figure 7:**
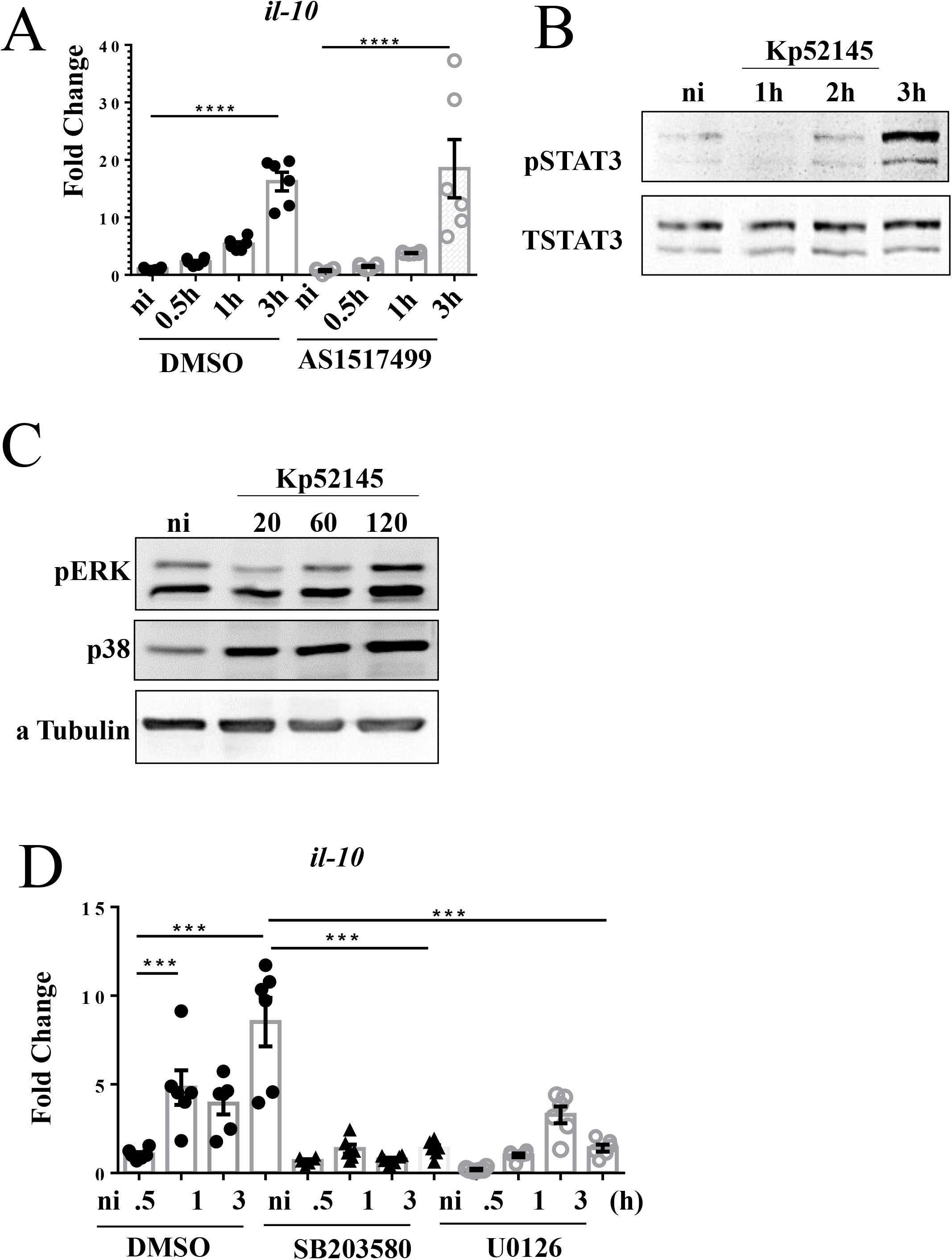
*K.pneumonia* induces *il-10* expression which is p38 and pERK–dependent. A. *il-10* levels in pBMDMs non-infected (ni) or infected with Kp52145 pre-treated with STAT6 inhibitor (AS 1517499, 50nM/ 2 h prior to infection) or DMSO vehicle control. Values are shown as standard error of mean of three independent experiments. B. Immunoblotting analysis of phosphorylation of STAT3 (PSTAT3) and STAT3 in lysates of pBMDMs infected with Kp52145 for the indicated times or left uninfected (ni). Data is representative of three independent experiments. C. Immunoblotting analysis of phosphorylations of ERK (pERK), p38 (Pp38), and tubulin in lysates of pBMDMs infected with Kp52145 for the indicated times or left uninfected (ni). Data is representative of three independent experiments. D. *il-10* levels in pBMDMs non-infected (ni) or infected with Kp52145 pre-treated with p38 inhibitor (SB203580, Tocris, 10 μg/mL, 2 h prior to infection), ERK inhibitor (U0126, LC laboratories, 20 μg/mL, 2 h prior to infection) or DMSO vehicle control. Values are shown as standard error of mean of three independent experiments. In panels A and D, ****p < 0.0001, ***p < 0.001, for the indicated comparisons using one-way ANOVA with Bonferroni correction.

Control experiments showed that Kp52145 infection induced the phosphorylation of p38 and ERK MAP kinases in pBMDMs (Figure 7C). As we anticipated, pharmacologic inhibition of p38 and ERK with SB203580 and U0126, respectively, resulted in decrease in the expression of *il-10* in infected pBMDMs (Figuere 7D).

Altogether, these results demonstrate that *K. pneumoniae* skews macrophage polarization towards a M2-state in an STAT6-dependent manner. Furthermore, our results indicate that *Klebsiella*-induced macrophage polarization is dependent on the CPS.

## DISCUSSION

The development of infection models that approximate human disease is essential not only for understanding pathogenesis at the molecular level, but also to test new therapies before entering into clinical stages. This is particularly relevant given the costs of clinical trials, and the impact on the health system. Animal models, chiefly rodents, have provided invaluable information and, not surprisingly, they are used in most of the pre-clinical studies. However, the limitations of these models in terms of yielding accurate pre-clinical data to inform clinical trials is widely recognized. Furthermore, in the context of infectious diseases it is an established fact the significant differences between rodents, mice, and humans in terms of immune activation following infection (2). In fact, the different immune/inflammatory pathways existing between mouse and man have fuelled the conclusion that murine studies are unreliable predictors of human outcomes. In this regard, porcine models are becoming increasingly important as ideal preclinical models. The anatomical and physiological similarities between pigs and humans, including the activation of the immune system, argue in favour of using pigs to model human diseases.

The results of this study strongly suggest that the EVLP model using pig lungs could be considered a platform to investigate the infection biology of respiratory pathogens and, eventually, to run pre-clinical studies testing new therapeutics. As a proof-of-principle, we provide evidence demonstrating that infection of the porcine EVLP with the human pathogen *K. pneumoniae* recapitulates the known features of *Klebsiella*-triggered pneumonia including the lung injury associated with the infection and the recruitment of neutrophils and other immune cells following infection. Moreover, our data revealed the EVLP model is useful to assess the virulence potential of *K. pneumoniae*. The *K. pneumoniae cps* mutant previously known to be attenuated in the mouse pneumonia model was also attenuated in the porcine EVLP model.

To set-up the EVLP infection model using pig lungs we took advantage of the advances in organ preparation for lung transplants. The EVLP method has become prevalent in lung transplant centres around the world (20), and has been proven as a mean to prolong the window for transplant evaluation (49). The lung transplant community has developed a robust protocol for EVLP that can capture key physiologic parameters (gas exchange, lung mechanics, pulmonary vascular hemodynamics, and oedema) and to obtain samples for limited analysis (20). In our study we have adapted the EVLP technology used for human lungs to pig lungs, and we have developed a robust infection method to assess the infection biology of respiratory pathogens. Although in this study we have focused on *K. pneumoniae*, the model is amenable to use with other bacterial pathogens, but also viruses and fungi.

*Ex vivo* modelling is superior to tissue- and cell-based assays because the architectural integrity of the lung is preserved. For example, type I pneumocytes, which cover over 90% of the gas exchange surface of the lung, are difficult to culture *in vitro*; therefore, little is known about the response of this cell type to injury and infection. Our model is a significant step change from the previous elegant infection model using ex vivo sections of pig lungs to assess bacterial virulence (50). This cell-free model allows to investigate pathogen physiology in a spatially structured environment. However, the porcine EVLP model developed here facilitates the study of the functional interactions between different immune cells, dendritic cells, neutrophils macrophages, and epithelial cells in a more physiological setting. The main advantage of using pig versus human lungs is the availability of the former. There is scarce number of human lungs not suitable for transplant, and the access to them is expensive. Nonetheless, several studies have proven that the EVLP model using human lungs is suitable to test disease-modifying therapies in acute lung injury to generate relevant, reliable and predictable human pharmacodynamic, pharmacokinetic and toxicology data through analysis at the organ (51). Recently, we have successfully adapted the EVLP model using human lungs to study *Klebsiella* infection biology.

Another novel finding of our study is that *Klebsiella* skews macrophage polarization to an M2-like state. Importantly, our findings uncovered that *Klebsiella*-induced macrophage polarization is dependent on the activation of STAT6, the most important transcriptional factor governing M2 polarization (43, 44). M1 phenotype is characterized by the expression of high levels of proinflammatory cytokines, high production of reactive oxygen intermediates and iNOS-dependent reactive nitrogen intermediates, promotion of Th1 response by IL12 production, and potent microbicidal activity (35, 42). In contrast, M2 macrophages are characterized by the selective expression of markers such as arginase 1 (Arg1), CD163 as well as the production of low levels of IL-12, iNOS, and enhanced IL-10 production (35, 42). M1 macrophages are generally considered responsible for resistance against intracellular pathogens (34). Not surprisingly, a growing number of studies show that some pathogens have evolved different strategies to interfere with M1 polarization (34) whereas there are few examples of intracellular pathogens (*Francisella*, *Salmonella*, *Coxiella*, *Tropheryma*) inducing an anti-inflammatory M2 state (52–55). The potential impact of *Klebsiella* on macrophage plasticity has been largely overlooked. Most likely this is due to the fact that *Klebsiella* has been traditionally considered an extracellular pathogen, although our laboratory has recently demonstrated that *Klebsiella* survives intracellular in mouse and human macrophages by preventing phagolysosome fusion (19). The facts that the attenuated *cps* mutant did not activate STAT6, and did not induce an M2-like state strongly suggest that the induction of an M2-like state is a virulence strategy of *Klebsiella* to promote infection. We and others have provided compelling evidence showing that *K. pneumoniae* CPS is a bona fide immune evasin (15, 18, 25, 56–59). The results of this study further reinforce this notion by demonstrating that the CPS skews macrophage polarizations towards an M2 state. Further studies are warranted to investigate whether this could be a general feature of other CPS.

Interestingly, our findings provide an explanation for the clinical observation that some health factors such as alcohol abuse or viral infections are associated with increased susceptibility to *Klebsiella* infections (60–62). These factors are known to increase the number of M2 macrophages in the lung (63–65) which then could facilitate *Klebsiella* infection. Supporting this hypothesis, there is an improvement in bacterial clearance when this macrophage population is eliminated *in vivo* (63–65).

Despite the clear utility of the EVLP model to assess infections, it is worthwhile commenting on the limitations. The process recapitulated in the EVLP model represent early steps in the infection process and do not model other aspects such as organ dissemination. In addition, the model does not integrate other signals, such as those from the gut, known to be relevant to control infections (66–68). Further impediments are the difficulties to generate cell specific knock-in or knock-outs and the relative low-throughput of the model to test several bacterial mutants. However, we believe that the advantages significantly outweighs the limitations, and the EVLP model is a useful translational pre-clinical model to illuminate new aspects of the infection biology of pathogens such as those identified in this work.

## MATERIAL AND METHODS

### Collection of lungs and whole blood

Immediately after euthanization, 200 mL of whole blood was collected rapidly in sterile receptacles containing 10% citrate phosphate dextrose solution (C7165, SIGMA), an anticoagulant, whole blood was then mixed gently and kept at room temperature. Both lungs and heart were promptly removed by sharp dissection. The heart was removed leaving ample (>3 cm) section of pulmonary artery intact. Lungs were then separated along the carina again leaving at least 3cm of trachea intact. As pigs have an additional bronchus (cranial) on the right lung, only left lungs were selected for this study as they are readily compatible with the LS1 system and are anatomically more similar to human lungs (Judge *et al.,* 2014). The left lung was then flushed gradually with 500 mL Dulbecco’s Modified Eagle Medium (DMEM) without phenol red via pulmonary artery to remove blood using a 50 mL syringe. Tissue was then wrapped in plastic and placed on ice for transportation to the lab. Lungs were rejected for this study if they contained large areas of haemorrhage or consolidation.

### Preparation of lungs for EVLP

A cannula was placed in pulmonary artery and connected to the efferent tube of the VivoLine LS1 reconditioning unit to facilitate perfusion. Similarly, an endobronchial tube was inserted into the bronchus and secured with suture before being connected to a ventilator circuit with adult bacterial viral filters (140141/1, DS Medical). The LS1 temperature probe was placed in pulmonary veins and secured in place using a surgical suture. “Perfusate” consisting of 2 L of DMEM (Invitrogen) without phenol red and supplemented with 5% L-glutamine and 5% fetal calf serum (FCS), was placed in the base of the reservoir. Target temperature was set to 37°C. Initial perfusion began with 0.05 L/min, at this point ensuring that the LS1 shunt is open, and flow gas gradually increased to 0.4 L/min maintaining a pulmonary artery pressure of 10 - 15 mmHg. Once a temperature of 30°C was reached, the lungs were gently inflated with an Ambu bag. Ensuring lung is warm prior to inflation reduces risk of capillary damage. Continuous positive airway pressure (CPAP) of 10 cm H_2_O was applied with 95%O_2_/5% CO_2_ using a mechanical ventilator (Dräger Evita). Once system reaches 36°C with desired pressure, 200 mL of autologous blood were added to the perfusate to act as a reservoir for immune cell recruitment.

### Bronchoalveolar lavage (BAL)

Once a temperature of 36°C was reached, a baseline broncho-alveolar lavage (BAL) sample was collected by inserting a catheter (PE 240-Harvard apparatus) into the sub-segment (caudal) lobe via the endotracheal tube and gently advanced until resistance was encountered, at which point the catheter was withdrawn by 1 cm. Then 125 mL of warmed normal saline was instilled and retrieved after 5 minutes through the same catheter. The catheter was then used to deliver 5 mL of sterile PBS or 5 × 10^5^ CFU Kp52145 in 5 mL of PBS. After 4 h BAL sampling was repeated prior to disconnecting the lung and tissue collection carried out. BAL samples were assessed for total protein and innate immune infiltrates.

### Bacterial preparation

*K. pneumoniae* 52.145, a clinical isolate (serotype O1:K2) previously described (22, 69) was utilised alongside the isogenic *cps* mutant, strain 52145-Δ*wca*_*K2*_, which has been previously described (70). Bacteria were tagged with GFP by transformation with plasmid pFPV25.1Cm (25). For infections, a single colony was cultured in 5 mL of LB broth overnight at 36°C with gentle agitation. After 1:10 dilution, bacteria were grown to exponential phase by incubation at 37°C with agitation for 2.5 h. Bacteria were then adjusted OD_600_ at 1.0 in PBS. For *in vitro* infections, macrophages were infected with a M.O.I of 100:1, whereas 5 × 10^5^ CFU/mL were used to infect the lungs. CFUs in the tissue were determined by homogenising 100 μg of tissue from caudal lobe in 1 mL sterile PBS and plating serial dilutions on *Salmonella-Shigella* -agar plates (SIGMA). Three samples were assessed across each lung. Plates were incubated overnight at 37°C before counting. When required, antibiotics were added to the growth medium at the following concentration: rifampicin, 50 µg/ml; chloramphenicol 25 µg/ml.

### Protein quantification

Protein concentration was assessed in BAL samples at baseline (0 h) and at 4 h. Standards and samples were incubated with Pierce 660 nm protein assay (150 µl of reagent: 10 µl of sample/standard) at room temperature for 5 min prior to quantification using the Nanodrop spectrophotometer as per manufacturer instructions (Thermo scientific).

### Histology

Tissue sections (~1 cm^3^) were collected from the cranial, middle and caudal lobes of each lung fixedplaced in 10% formalin × 10 volume of tissue sample in a 15 mL Falcon tube with an inverted p1000 tip to submerge tissue. After a minimum of 48 h at room temperature, samples were then processed for paraffin embedding, sectioning and haematoxylin and eosin staining. Samples were imaged using the DM5500 Leica vertical microscope at × 200 magnification. Alveolar septal oedema was quantified by measuring alveolar septal thickness with ImageJ software, whereby three measurements of thickest septa were acquired per image and averaged, 30 images were acquired whereby 10 images were acquired per section and 3 sections per lung.. Alveolar septa adjacent to blood vessel or airway were excluded due to normal thickening resulting from collagen deposition. Intra-alveolar hemorrhage, presence of intra-alveolar mononuclear cells and proteinaceous debris were also recorded. Histological scores were assigned based on parameters set by Matute-Bello and co-workers (30). Whereby hemorrhage was scored as follows 0= none, 1= mild, 2= moderate and 3= severe. Proteinaceous debris scored as 0 = none, 1 = protein present and 2= abundant presence of protein in alveolar spaces. The number of nucleated cells within the alveolar space were counted and presented as intra-alveolar leukocytes. 5 images were scored per section with 3 sections per lung at × 400 magnification.

### RNA purifcation

100 μg of lung tissue was homogenised using a VDI 12 tissue homogeniser (VWR) in 1 mL of TRizol reagent (Ambion) and incubated at room temperature for 5 min before storing at −80 °C. RNA was extracted from pBMDMs using RNeasy^®^ Minikit (QIAGEN ref: 74104). Total RNA was extracted according to manufacturer’s instructions. 5 μg of total RNA were treated with recombinant DNase I (Roche Diagnostics Ltd.) at 37°C for 30 min and then purified using a standard phenol–chloroform method. The RNA was precipitated with 20 μl 3 M sodium acetate (pH 5.2) and 600 μ1 98% (v/v) ethanol at −20°C, washed twice in 75% (v/v) ethanol, dried and then resuspended in RNase-free H2O. Duplicate cDNA preparations from each sample were generated from 1 μg of RNA using Moloney murine leukaemia virus (M-MLV) reverse transcriptase (Sigma-Aldrich) according to the manufacturer’s instructions. RT-qPCR analysis of cytokine related porcine gene expression was performed using the KAPA SYBR-FAST qPCR Kit (KAPA Biosystems), using the primers shown in Table 1. Samples were run using the Stratagene Mx3005P qPCR System (Agilent Technologies). Non-template negative controls to check for primer-dimer and a porcine genomic DNA were included. Thermal cycling conditions were as follows: 95°C for 3 min for enzyme activation, 40 cycles of denaturation at 95°C for 10 s and annealing at 60°C for 20 s. cDNA samples were tested in duplicate and relative mRNA quantity was determined by the comparative threshold cycle (ΔΔCt) method using HPRT housekeeping gene for normalisation.

**Table 1.**
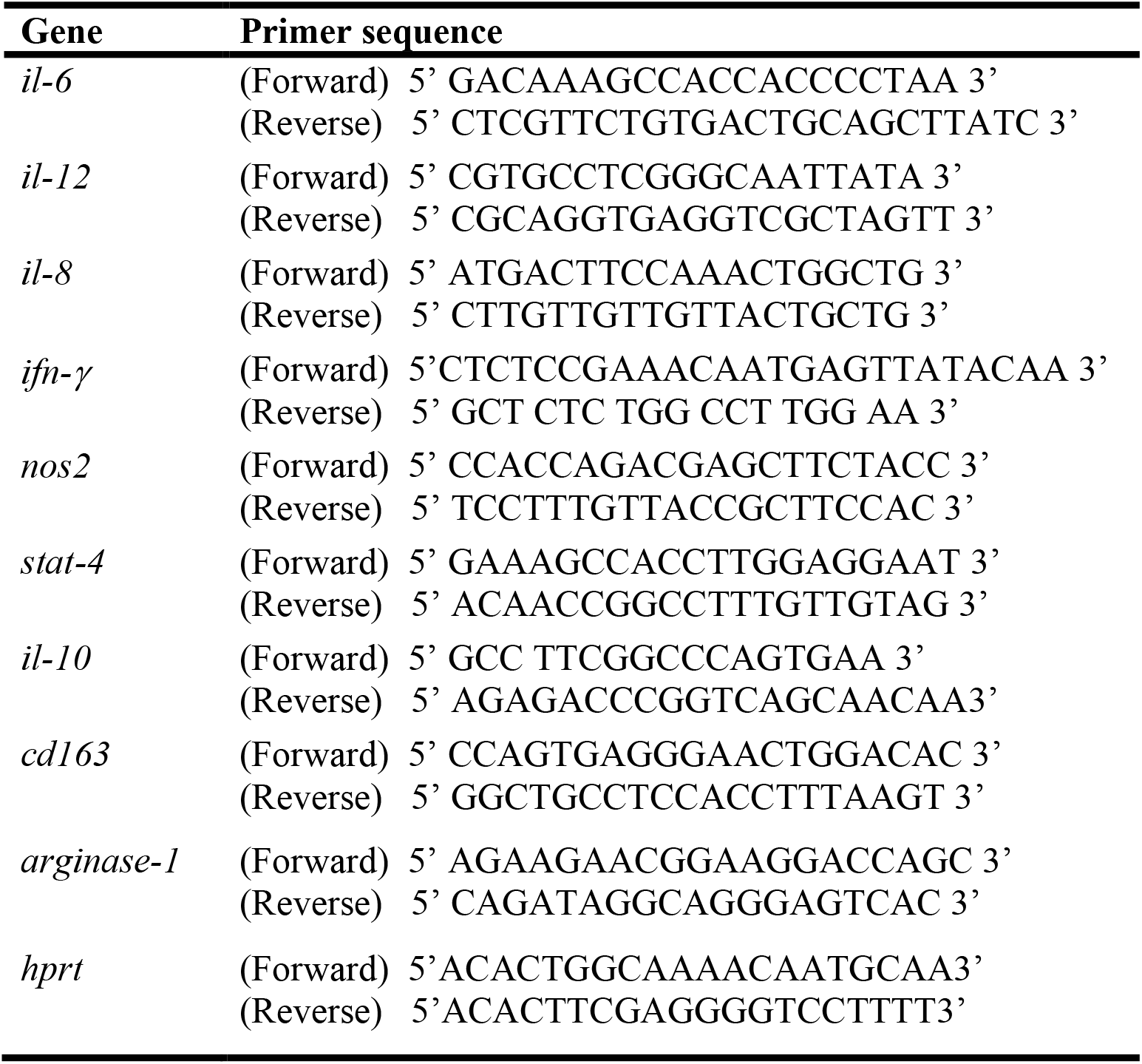
List of primers used in this study for RT-qPCR.

### Flow cytometry

100 μg of lung tissue was homogenised in 1 mL sterile PBS and filtered through a 70 μm cell strainer (2236348 Fisherbrand). Cells were centrifuged and red cells lysed using ammonium-chloride-potassium lysis buffer (A1049201, Gibco) for 3 min at room temperature, washed with 1 mL PBS prior to staining with the following mouse anti-pig antibodies: CD11R3 (MCA2309), CD163 (clone 2A10), SLA Class II and Granulocyte antibody (clone 6D10) (AbD Serotech). Each purified anti-pig antibody was labelled with a fluorophore using Abcam conjugation kits PE (ab102918), APC-Cy5.5 (ab102855), FITC (ab102884), and Rhodamine (ab188286)

### Generation of porcine bone marrow-derived macrophages (pBMDMs)

Femurs from pigs between 80-100 kg were cleared of all muscle and sinew. Bone was then washed with 70 % ethanol. A sterilised junior hacksaw was used to cut transversely across bone to expose bone marrow under sterile conditions. 5 g of bone marrow per 50 mL tube were suspended in 40 mL complete media and centrifuged at 600 × g for 8 min to remove fat. Red cells were lysed via incubations with ammonium-chloride-potassium lysis buffer (A1049201, Gibco) for 3 min. Cells were washed in 10 mL complete media and passed through a 70 mm cell strainer (2236348 Fisherbrand) prior to centrifugation. Cell pellet was dislodged before plating on 20 cm petri dishes (SARSTEDT) in 25 mL complete medium (DMEM, high glucose, GlutaMAX™, supplemented with 10% FCS, 1% pen/strep) and 5 mL of syringe filtered L929 supernatant (a source of M-CSF). Cells were cultured for 6 days before assessment of purity by flow cytometry.

### *In vitro* infections

pBMDMs were seeded in 6 well dishes (5 × 10^5^ cells/well) in complete media (DMEM, high glucose, GlutaMAX™, supplemented with 10% FCS and 1% pen/strep) and allowed to adhere overnight. Complete media was removed and replaced with antibiotic free media prior to infection. Bacterial inoculum was prepared as previously indicated and cells were infected with a M.O.I of 100 bacteria per cell. To synchronise infection, plates were centrifuged at 200 × g for 5 min. After 1 h, media was removed, replaced with antibiotic free media supplemented with 100 µg/ml gentamicin (SIGMA) to kill extracellular bacteria. For STAT6 inhibition, cells were serum starved and incubated with the chemical STAT6 inhibitor AS 1517499 (50 nM, 919486-40-1, AXON Medchem) or DMSO as vehicle control for 2 h prior to infection and maintained throughout. To inhibit pERK and p38 activity, the chemical inhibitors U0126 (20 μg/mL, LC laboratories,) and SB203580 (10 μg/mL, Tocris,) were utilised respectively 2 h prior to infection and maintained throughout experiment. At indicated time points, supernatants were removed and cells lysed for analysis by western blotting or qRT-PCR.

### Western blotting

At appropriate time point post-infection, cells were washed with ice-cold PBS before lysis in Laemmli buffer (4% SDS, 10% 2-mercaptoehtanol, 20% glycerol, 0.004% bromophenol blue, 0.125 M Tris-HCl pH 6.8). Lysates were sonicated for 10 sec at 10% amplitude, boiled at 95 ºC for 5 minutes and centrifuged at 12,000g for 1 min prior to running on 8% SDS-PAGE. Samples were transferred onto 0.2 mm nitrocellulose membrane (Biotrace, VWR) using a semi-dry transfer unit (Bio-Rad) before blocking nonspecific antibody binding for 1 h in 3% BSA in TBS with 1 % Tween-20. Primary antibodies included: phospho-STAT6 (Tyr641) (1:2000, #9361), total STAT6 (1:1000, BioRad #170-6516), phospho-STAT3 (Y705) (1:2000, #9145), total STAT3 (1:2000, #12640), phospho-ERK (p44/42) (1:2000; #91015), phopsho-p38 (T180/Y182) (1:2000, #4511), all from Cell Signalling Technologies. Total STAT6 (21HCLC) (1:1000, Thermo Scientific, #701110). Blots were incubated with appropriate horseradish peroxidase -conjugated secondary antibody goat anti-rabbit immunoglobulins (1:5000, BioRad 170-6515) or goat anti-mouse immunoglobulins (1:1000, BioRad 170-6516). Protein bands were detected using chemiluminescence reagents and a G:BOX Chemi XRQ chemiluminescence imager (Syngene). To detect multiple proteins, membranes were reprobed after stripping of previously used antibodies using a pH 2.2 glycine-HCl/SDS buffer. To ensure that equal amounts of proteins were loaded, blots were reprobed with α-tubulin (1:2000, Cell Signalling Technologies #2125).

### Statistics

Statistical analyses were performed with Prism 6 (GraphPad Software) using 1-way ANOVA with Bonferroni correction or unpaired two-tailed Student’s t-test. Error bars indicate standard error of mean (SEM) Statistical significance is indicated as follows: ns (not significant), P > 0.05, *, P < 0.05; **, P < 0.01; ***, P < 0.001.

## ACKNOWLEDGEMENTS

We thank the members of the J.A.B. laboratory for their thoughtful discussions and support with this project. This work was supported by Marie Curie Career Integration Grant U-KARE (PCIG13-GA-2013-618162); Biotechnology and Biological Sciences Research Council (BBSRC, BB/P006078/1), Medical Research Council (MR/R005893/1) and Queen’s University Belfast start-up funds to J.A.B.

## AUTHORS CONTRIBUTIONS

AD, DFM, CMO and JAB conceived the study and wrote the first draft of the manuscript. AD, JS-P, MF, UH performed the experiments and contributed data for this work. AD, MF, UH, JS-P, DFM, CMO, and JAB contributed to and approved the final version of the manuscript.

## Supplementary Figure Legends

**Supplementary figure 1: Regions for sample selection from porcine EVLP lung**.

Image identifying “cranial”, “middle” and “caudal” regions for tissue sample collection.

**Supplementary figure 2: Innate cells recruitment in *K. pneumoniae*-infected EVLP porcine model**.

Gating strategy and representative dot plots for flow cytometric analysis of A) CD11R3+ macrophages, B) Neutrophil staining using anti-pig granulocyte marker clone 6D10, C) Analysis of CD11R3-CD172+ dendritic cells, Dot plots represent 0h (baseline) and 4h post-infection or mock infection BAL samples and 4h tissue samples. D) Gating strategy to identify differential expression of M2 marker CD163 on K.p infected macrophages (CD11R3+GFP+CD163+) or K.p (-) macrophages (CD11R3+GFP-CD163+).

**Supplementary figure 3: *K. pneumoniae* CPS mutant induces a M1 markers in pBMDMs**.

*Stat4* and *nos2* levels in pBMDMs non-infected (ni) or infected with *cps* mutant, strain 52145-Δ*wca*_*K2*_, pre-treated with STAT6 inhibitor (AS1517499, 50nM 2 h prior to infection) or DMSO vehicle control. Values are shown as standard error of mean of three independent experiments in duplicate. ****p<0.0001, **p<0.01, n.s. p>0.05 for the indicated comparisons using one-way ANOVA with Bonferroni correction.

## References

1 Greek R, Menache A. 2013. Systematic reviews of animal models: methodology versus epistemology. Int J Med Sci 10:206–221.

2 Mizgerd JP, Skerrett SJ. 2008. Animal models of human pneumonia. Am J Physiol Lung Cell Mol Physiol 294:L387–98.

3 Mouse Genome Sequencing Consortium, Waterston RH, Lindblad-Toh K, Birney E, Rogers J, Abril JF, Agarwal P, Agarwala R, Ainscough R, Alexandersson M, An P, Antonarakis SE, Attwood J, Baertsch R, Bailey J, Barlow K, Beck S, Berry E, Birren B, Bloom T, Bork P, Botcherby M, Bray N, Brent MR, Brown DG, Brown SD, Bult C, Burton J, Butler J, Campbell RD, Carninci P, Cawley S, Chiaromonte F, Chinwalla AT, Church DM, Clamp M, Clee C, Collins FS, Cook LL, Copley RR, Coulson A, Couronne O, Cuff J, Curwen V, Cutts T, Daly M, David R, Davies J, Delehaunty KD, Deri J, Dermitzakis ET, Dewey C, Dickens NJ, Diekhans M, Dodge S, Dubchak I, Dunn DM, Eddy SR, Elnitski L, Emes RD, Eswara P, Eyras E, Felsenfeld A, Fewell GA, Flicek P, Foley K, Frankel WN, Fulton LA, Fulton RS, Furey TS, Gage D, Gibbs RA, Glusman G, Gnerre S, Goldman N, Goodstadt L, Grafham D, Graves TA, Green ED, Gregory S, Guigo R, Guyer M, Hardison RC, Haussler D, Hayashizaki Y, Hillier LW, Hinrichs A, Hlavina W, Holzer T, Hsu F, Hua A, Hubbard T, Hunt A, Jackson I, Jaffe DB, Johnson LS, Jones M, Jones TA, Joy A, Kamal M, Karlsson EK, Karolchik D, Kasprzyk A, Kawai J, Keibler E, Kells C, Kent WJ, Kirby A, Kolbe DL, Korf I, Kucherlapati RS, Kulbokas EJ, Kulp D, Landers T, Leger JP, Leonard S, Letunic I, Levine R, Li J, Li M, Lloyd C, Lucas S, Ma B, Maglott DR, Mardis ER, Matthews L, Mauceli E, Mayer JH, McCarthy M, McCombie WR, McLaren S, McLay K, McPherson JD, Meldrim J, Meredith B, Mesirov JP, Miller W, Miner TL, Mongin E, Montgomery KT, Morgan M, Mott R, Mullikin JC, Muzny DM, Nash WE, Nelson JO, Nhan MN, Nicol R, Ning Z, Nusbaum C, O’Connor MJ, Okazaki Y, Oliver K, Overton-Larty E, Pachter L, Parra G, Pepin KH, Peterson J, Pevzner P, Plumb R, Pohl CS, Poliakov A, Ponce TC, Ponting CP, Potter S, Quail M, Reymond A, Roe BA, Roskin KM, Rubin EM, Rust AG, Santos R, Sapojnikov V, Schultz B, Schultz J, Schwartz MS, Schwartz S, Scott C, Seaman S, Searle S, Sharpe T, Sheridan A, Shownkeen R, Sims S, Singer JB, Slater G, Smit A, Smith DR, Spencer B, Stabenau A, Stange-Thomann N, Sugnet C, Suyama M, Tesler G, Thompson J, Torrents D, Trevaskis E, Tromp J, Ucla C, Ureta-Vidal A, Vinson JP, Von Niederhausern AC, Wade CM, Wall M, Weber RJ, Weiss RB, Wendl MC, West AP, Wetterstrand K, Wheeler R, Whelan S, Wierzbowski J, Willey D, Williams S, Wilson RK, Winter E, Worley KC, Wyman D, Yang S, Yang SP, Zdobnov EM, Zody MC, Lander ES. 2002. Initial sequencing and comparative analysis of the mouse genome. Nature 420:520–562.

4 Glavis-Bloom J, Muhammed M, Mylonakis E. 2012. Of model hosts and man: using Caenorhabditis elegans, Drosophila melanogaster and Galleria mellonella as model hosts for infectious disease research. Adv Exp Med Biol 710:11–17.

5 Torraca V, Mostowy S. 2018. Zebrafish Infection: From Pathogenesis to Cell Biology. Trends Cell Biol 28:143–156.

6 Groenen MA, Archibald AL, Uenishi H, Tuggle CK, Takeuchi Y, Rothschild MF, Rogel-Gaillard C, Park C, Milan D, Megens HJ, Li S, Larkin DM, Kim H, Frantz LA, Caccamo M, Ahn H, Aken BL, Anselmo A, Anthon C, Auvil L, Badaoui B, Beattie CW, Bendixen C, Berman D, Blecha F, Blomberg J, Bolund L, Bosse M, Botti S, Bujie Z, Bystrom M, Capitanu B, Carvalho-Silva D, Chardon P, Chen C, Cheng R, Choi SH, Chow W, Clark RC, Clee C, Crooijmans RP, Dawson HD, Dehais P, De Sapio F, Dibbits B, Drou N, Du ZQ, Eversole K, Fadista J, Fairley S, Faraut T, Faulkner GJ, Fowler KE, Fredholm M, Fritz E, Gilbert JG, Giuffra E, Gorodkin J, Griffin DK, Harrow JL, Hayward A, Howe K, Hu ZL, Humphray SJ, Hunt T, Hornshoj H, Jeon JT, Jern P, Jones M, Jurka J, Kanamori H, Kapetanovic R, Kim J, Kim JH, Kim KW, Kim TH, Larson G, Lee K, Lee KT, Leggett R, Lewin HA, Li Y, Liu W, Loveland JE, Lu Y, Lunney JK, Ma J, Madsen O, Mann K, Matthews L, McLaren S, Morozumi T, Murtaugh MP, Narayan J, Nguyen DT, Ni P, Oh SJ, Onteru S, Panitz F, Park EW, Park HS, Pascal G, Paudel Y, Perez-Enciso M, Ramirez-Gonzalez R, Reecy JM, Rodriguez-Zas S, Rohrer GA, Rund L, Sang Y, Schachtschneider K, Schraiber JG, Schwartz J, Scobie L, Scott C, Searle S, Servin B, Southey BR, Sperber G, Stadler P, Sweedler JV, Tafer H, Thomsen B, Wali R, Wang J, Wang J, White S, Xu X, Yerle M, Zhang G, Zhang J, Zhang J, Zhao S, Rogers J, Churcher C, Schook LB. 2012. Analyses of pig genomes provide insight into porcine demography and evolution. Nature 491:393–398.

7 Aigner B, Renner S, Kessler B, Klymiuk N, Kurome M, Wunsch A, Wolf E. 2010. Transgenic pigs as models for translational biomedical research. J Mol Med (Berl) 88:653–664.

8 Munoz-Price LS, Poirel L, Bonomo RA, Schwaber MJ, Daikos GL, Cormican M, Cornaglia G, Garau J, Gniadkowski M, Hayden MK, Kumarasamy K, Livermore DM, Maya JJ, Nordmann P, Patel JB, Paterson DL, Pitout J, Villegas MV, Wang H, Woodford N, Quinn JP. 2013. Clinical epidemiology of the global expansion of Klebsiella pneumoniae carbapenemases. Lancet Infect Dis 13:785–796.

9 Bengoechea JA, Sa Pessoa J. 2019. Klebsiella pneumoniae infection biology: living to counteract host defences. FEMS Microbiol Rev 43:123–144.

10 Ivin M, Dumigan A, de Vasconcelos FN, Ebner F, Borroni M, Kavirayani A, Przybyszewska KN, Ingram RJ, Lienenklaus S, Kalinke U, Stoiber D, Bengoechea JA, Kovarik P. 2017. Natural killer cell-intrinsic type I IFN signaling controls Klebsiella pneumoniae growth during lung infection. PLoS Pathog 13:e1006696.

11 Xiong H, Keith JW, Samilo DW, Carter RA, Leiner IM, Pamer EG. 2016. Innate Lymphocyte/Ly6C(hi) Monocyte Crosstalk Promotes Klebsiella pneumoniae Clearance. Cell 165:679–689.

12 Xiong H, Carter RA, Leiner IM, Tang YW, Chen L, Kreiswirth BN, Pamer EG. 2015. Distinct Contributions of Neutrophils and CCR2+ Monocytes to Pulmonary Clearance of Different Klebsiella pneumoniae Strains. Infect Immun 83:3418–3427.

13 Broug-Holub E, Toews GB, van Iwaarden JF, Strieter RM, Kunkel SL, Paine R,3rd, Standiford TJ. 1997. Alveolar macrophages are required for protective pulmonary defenses in murine Klebsiella pneumonia: elimination of alveolar macrophages increases neutrophil recruitment but decreases bacterial clearance and survival. Infect Immun 65:1139–1146.

14 Cheung DO, Halsey K, Speert DP. 2000. Role of pulmonary alveolar macrophages in defense of the lung against Pseudomonas aeruginosa. Infect Immun 68:4585–4592.

15 Tomas A, Lery L, Regueiro V, Perez-Gutierrez C, Martinez V, Moranta D, Llobet E, Gonzalez-Nicolau M, Insua JL, Tomas JM, Sansonetti PJ, Tournebize R, Bengoechea JA. 2015. Functional Genomic Screen Identifies Klebsiella pneumoniae Factors Implicated in Blocking Nuclear Factor kappaB (NF-kappaB) Signaling. J Biol Chem 290:16678–16697.

16 March C, Moranta D, Regueiro V, Llobet E, Tomas A, Garmendia J, Bengoechea JA. 2011. Klebsiella pneumoniae outer membrane protein A is required to prevent the activation of airway epithelial cells. J Biol Chem 286:9956–9967.

17 Llobet E, Martinez-Moliner V, Moranta D, Dahlstrom KM, Regueiro V, Tomas A, Cano V, Perez-Gutierrez C, Frank CG, Fernandez-Carrasco H, Insua JL, Salminen TA, Garmendia J, Bengoechea JA. 2015. Deciphering tissue-induced Klebsiella pneumoniae lipid A structure. Proc Natl Acad Sci U S A 112:E6369–E6378.

18 Lawlor MS, Handley SA, Miller VL. 2006. Comparison of the host responses to wild-type and cpsB mutant Klebsiella pneumoniae infections. Infect Immun 74:5402–5407.

19 Cano V, March C, Insua JL, Aguilo N, Llobet E, Moranta D, Regueiro V, Brennan GP, Millan-Lou MI, Martin C, Garmendia J, Bengoechea JA. 2015. Klebsiella pneumoniae survives within macrophages by avoiding delivery to lysosomes. Cell Microbiol 17:1537–1560.

20 Tane S, Noda K, Shigemura N. 2017. Ex Vivo Lung Perfusion: A Key Tool for Translational Science in the Lungs. Chest 151:1220–1228.

21 Judge EP, Hughes JM, Egan JJ, Maguire M, Molloy EL, O’Dea S. 2014. Anatomy and bronchoscopy of the porcine lung. A model for translational respiratory medicine. Am J Respir Cell Mol Biol 51:334–343.

22 Lery LM, Frangeul L, Tomas A, Passet V, Almeida AS, Bialek-Davenet S, Barbe V, Bengoechea JA, Sansonetti P, Brisse S, Tournebize R. 2014. Comparative analysis of Klebsiella pneumoniae genomes identifies a phospholipase D family protein as a novel virulence factor. BMC Biol 12:41-7007-12-41.

23 Holt KE, Wertheim H, Zadoks RN, Baker S, Whitehouse CA, Dance D, Jenney A, Connor TR, Hsu LY, Severin J, Brisse S, Cao H, Wilksch J, Gorrie C, Schultz MB, Edwards DJ, Nguyen KV, Nguyen TV, Dao TT, Mensink M, Minh VL, Nhu NT, Schultsz C, Kuntaman K, Newton PN, Moore CE, Strugnell RA, Thomson NR. 2015. Genomic analysis of diversity, population structure, virulence, and antimicrobial resistance in Klebsiella pneumoniae, an urgent threat to public health. Proc Natl Acad Sci U S A 112:E3574–E3581.

24 Insua JL, Llobet E, Moranta D, Perez-Gutierrez C, Tomas A, Garmendia J, Bengoechea JA. 2013. Modeling Klebsiella pneumoniae pathogenesis by infection of the wax moth Galleria mellonella. Infect Immun 81:3552–3565.

25 March C, Cano V, Moranta D, Llobet E, Perez-Gutierrez C, Tomas JM, Suarez T, Garmendia J, Bengoechea JA. 2013. Role of bacterial surface structures on the interaction of Klebsiella pneumoniae with phagocytes. PLoS One 8:e56847.

26 Tournebize R, Doan BT, Dillies MA, Maurin S, Beloeil JC, Sansonetti PJ. 2006. Magnetic resonance imaging of Klebsiella pneumoniae-induced pneumonia in mice. Cell Microbiol 8:33–43.

27 Izquierdo L, Coderch N, Pique N, Bedini E, Corsaro MM, Merino S, Fresno S, Tomas JM, Regue M. 2003. The Klebsiella pneumoniae wabG gene: role in biosynthesis of the core lipopolysaccharide and virulence. J Bacteriol 185:7213–7221.

28 Cortes G, Borrell N, de Astorza B, Gomez C, Sauleda J, Alberti S. 2002. Molecular analysis of the contribution of the capsular polysaccharide and the lipopolysaccharide O side chain to the virulence of Klebsiella pneumoniae in a murine model of pneumonia. Infect Immun 70:2583–2590.

29 Lawlor MS, Hsu J, Rick PD, Miller VL. 2005. Identification of Klebsiella pneumoniae virulence determinants using an intranasal infection model. Mol Microbiol 58:1054–1073.

30 Matute-Bello G, Downey G, Moore BB, Groshong SD, Matthay MA, Slutsky AS, Kuebler WM, Acute Lung Injury in Animals Study Group. 2011. An official American Thoracic Society workshop report: features and measurements of experimental acute lung injury in animals. Am J Respir Cell Mol Biol 44:725–738.

31 Summerfield A, Haverson K, Thacker E, McCullough KC. 2001. Differentiation of porcine myeloid bone marrow haematopoietic cell populations. Vet Immunol Immunopathol 80:121–129.

32 Thacker E, Summerfield A, McCullough K, Ezquerra A, Dominguez J, Alonso F, Lunney J, Sinkora J, Haverson K. 2001. Summary of workshop findings for porcine myelomonocytic markers. Vet Immunol Immunopathol 80:93–109.

33 Facci MR, Auray G, Buchanan R, van Kessel J, Thompson DR, Mackenzie-Dyck S, Babiuk LA, Gerdts V. 2010. A comparison between isolated blood dendritic cells and monocyte-derived dendritic cells in pigs. Immunology 129:396–405.

34 Benoit M, Desnues B, Mege JL. 2008. Macrophage polarization in bacterial infections. J Immunol 181:3733–3739.

35 Xue J, Schmidt SV, Sander J, Draffehn A, Krebs W, Quester I, De Nardo D, Gohel TD, Emde M, Schmidleithner L, Ganesan H, Nino-Castro A, Mallmann MR, Labzin L, Theis H, Kraut M, Beyer M, Latz E, Freeman TC, Ulas T, Schultze JL. 2014. Transcriptome-based network analysis reveals a spectrum model of human macrophage activation. Immunity 40:274–288.

36 Yoshida K, Matsumoto T, Tateda K, Uchida K, Tsujimoto S, Iwakurai Y, Yamaguchi K. 2001. Protection against pulmonary infection with Klebsiella pneumoniae in mice by interferon-gamma through activation of phagocytic cells and stimulation of production of other cytokines. J Med Microbiol 50:959–964.

37 Moore TA, Perry ML, Getsoian AG, Newstead MW, Standiford TJ. 2002. Divergent role of gamma interferon in a murine model of pulmonary versus systemic Klebsiella pneumoniae infection. Infect Immun 70:6310–6318.

38 Zeng X, Moore TA, Newstead MW, Deng JC, Kunkel SL, Luster AD, Standiford TJ. 2005. Interferon-inducible protein 10, but not monokine induced by gamma interferon, promotes protective type 1 immunity in murine Klebsiella pneumoniae pneumonia. Infect Immun 73:8226–8236.

39 Zeng X, Moore TA, Newstead MW, Deng JC, Lukacs NW, Standiford TJ. 2005. IP-10 mediates selective mononuclear cell accumulation and activation in response to intrapulmonary transgenic expression and during adenovirus-induced pulmonary inflammation. J Interferon Cytokine Res 25:103–112.

40 Yoshida K, Matsumoto T, Tateda K, Uchida K, Tsujimoto S, Yamaguchi K. 2001. Induction of interleukin-10 and down-regulation of cytokine production by Klebsiella pneumoniae capsule in mice with pulmonary infection. J Med Microbiol 50:456–461.

41 Yoshida K, Matsumoto T, Tateda K, Uchida K, Tsujimoto S, Yamaguchi K. 2000. Role of bacterial capsule in local and systemic inflammatory responses of mice during pulmonary infection with Klebsiella pneumoniae. J Med Microbiol 49:1003–1010.

42 Murray PJ, Allen JE, Biswas SK, Fisher EA, Gilroy DW, Goerdt S, Gordon S, Hamilton JA, Ivashkiv LB, Lawrence T, Locati M, Mantovani A, Martinez FO, Mege JL, Mosser DM, Natoli G, Saeij JP, Schultze JL, Shirey KA, Sica A, Suttles J, Udalova I, van Ginderachter JA, Vogel SN, Wynn TA. 2014. Macrophage activation and polarization: nomenclature and experimental guidelines. Immunity 41:14–20.

43 Liao X, Sharma N, Kapadia F, Zhou G, Lu Y, Hong H, Paruchuri K, Mahabeleshwar GH, Dalmas E, Venteclef N, Flask CA, Kim J, Doreian BW, Lu KQ, Kaestner KH, Hamik A, Clement K, Jain MK. 2011. Kruppel-like factor 4 regulates macrophage polarization. J Clin Invest 121:2736–2749.

44 Biswas SK, Mantovani A. 2012. Orchestration of metabolism by macrophages. Cell Metab 15:432–437.

45 Mikita T, Campbell D, Wu P, Williamson K, Schindler U. 1996. Requirements for interleukin-4-induced gene expression and functional characterization of Stat6. Mol Cell Biol 16:5811–5820.

46 Mikita T, Daniel C, Wu P, Schindler U. 1998. Mutational analysis of the STAT6 SH2 domain. J Biol Chem 273:17634–17642.

47 Nagashima S, Yokota M, Nakai E, Kuromitsu S, Ohga K, Takeuchi M, Tsukamoto S, Ohta M. 2007. Synthesis and evaluation of 2-{[2-(4-hydroxyphenyl)-ethyl]amino}pyrimidine-5-carboxamide derivatives as novel STAT6 inhibitors. Bioorg Med Chem 15:1044–1055.

48 Saraiva M, O’Garra A. 2010. The regulation of IL-10 production by immune cells. Nat Rev Immunol 10:170–181.

49 Cypel M, Yeung JC, Liu M, Anraku M, Chen F, Karolak W, Sato M, Laratta J, Azad S, Madonik M, Chow CW, Chaparro C, Hutcheon M, Singer LG, Slutsky AS, Yasufuku K, de Perrot M, Pierre AF, Waddell TK, Keshavjee S. 2011. Normothermic ex vivo lung perfusion in clinical lung transplantation. N Engl J Med 364:1431–1440.

50 Harrison F, Muruli A, Higgins S, Diggle SP. 2014. Development of an ex vivo porcine lung model for studying growth, virulence, and signaling of Pseudomonas aeruginosa. Infect Immun 82:3312–3323.

51 Proudfoot AG, McAuley DF, Griffiths MJ, Hind M. 2011. Human models of acute lung injury. Dis Model Mech 4:145–153.

52 Benoit M, Barbarat B, Bernard A, Olive D, Mege JL. 2008. Coxiella burnetii, the agent of Q fever, stimulates an atypical M2 activation program in human macrophages. Eur J Immunol 38:1065–1070.

53 Desnues B, Lepidi H, Raoult D, Mege JL. 2005. Whipple disease: intestinal infiltrating cells exhibit a transcriptional pattern of M2/alternatively activated macrophages. J Infect Dis 192:1642–1646.

54 Eisele NA, Ruby T, Jacobson A, Manzanillo PS, Cox JS, Lam L, Mukundan L, Chawla A, Monack DM. 2013. Salmonella require the fatty acid regulator PPARdelta for the establishment of a metabolic environment essential for long-term persistence. Cell Host Microbe 14:171–182.

55 Saliba AE, Li L, Westermann AJ, Appenzeller S, Stapels DA, Schulte LN, Helaine S, Vogel J. 2016. Single-cell RNA-seq ties macrophage polarization to growth rate of intracellular Salmonella. Nat Microbiol 2:16206.

56 Regueiro V, Campos MA, Pons J, Alberti S, Bengoechea JA. 2006. The uptake of a Klebsiella pneumoniae capsule polysaccharide mutant triggers an inflammatory response by human airway epithelial cellsy4. Microbiology 152:555–566.

57 Campos MA, Vargas MA, Regueiro V, Llompart CM, Alberti S, Bengoechea JA. 2004. Capsule polysaccharide mediates bacterial resistance to antimicrobial peptides. Infect Immun 72:7107–7114. 10.1128/IAI.72.12.7107-7114.2004.

58 Moranta D, Regueiro V, March C, Llobet E, Margareto J, Larrarte E, Garmendia J, Bengoechea JA. 2010. Klebsiella pneumoniae capsule polysaccharide impedes the expression of beta-defensins by airway epithelial cells. Infect Immun 78:1135–1146.

59 Frank CG, Reguerio V, Rother M, Moranta D, Maeurer AP, Garmendia J, Meyer TF, Bengoechea JA. 2013. Klebsiella pneumoniae targets an EGF receptor-dependent pathway to subvert inflammation. Cell Microbiol 15:1212–1233.

60 Ballinger MN, Standiford TJ. 2010. Postinfluenza bacterial pneumonia: host defenses gone awry. J Interferon Cytokine Res 30:643–652.

61 Happel KI, Odden AR, Zhang P, Shellito JE, Bagby GJ, Nelson S. 2006. Acute alcohol intoxication suppresses the interleukin 23 response to Klebsiella pneumoniae infection. Alcohol Clin Exp Res 30:1200–1207.

62 Mancuso P, Gottschalk A, Phare SM, Peters-Golden M, Lukacs NW, Huffnagle GB. 2002. Leptin-deficient mice exhibit impaired host defense in Gram-negative pneumonia. J Immunol 168:4018–4024.

63 Tsuchimoto Y, Asai A, Tsuda Y, Ito I, Nishiguchi T, Garcia MC, Suzuki S, Kobayashi M, Higuchi K, Suzuki F. 2015. M2b Monocytes Provoke Bacterial Pneumonia and Gut Bacteria-Associated Sepsis in Alcoholics. J Immunol 195:5169–5177.

64 Ohama H, Asai A, Ito I, Suzuki S, Kobayashi M, Higuchi K, Suzuki F. 2015. M2b macrophage elimination and improved resistance of mice with chronic alcohol consumption to opportunistic infections. Am J Pathol 185:420–431.

65 Dolgachev VA, Yu B, Reinke JM, Raghavendran K, Hemmila MR. 2012. Host susceptibility to gram-negative pneumonia after lung contusion. J Trauma Acute Care Surg 72:614-22; discussion 622–3.

66 Fagundes CT, Amaral FA, Vieira AT, Soares AC, Pinho V, Nicoli JR, Vieira LQ, Teixeira MM, Souza DG. 2012. Transient TLR activation restores inflammatory response and ability to control pulmonary bacterial infection in germfree mice. J Immunol 188:1411–1420.

67 Brown RL, Sequeira RP, Clarke TB. 2017. The microbiota protects against respiratory infection via GM-CSF signaling. Nat Commun 8:1512-017-01803-x.

68 Clarke TB. 2014. Early innate immunity to bacterial infection in the lung is regulated systemically by the commensal microbiota via nod-like receptor ligands. Infect Immun 82:4596–4606.

69 Nassif X, Sansonetti PJ. 1986. Correlation of the virulence of Klebsiella pneumoniae K1 and K2 with the presence of a plasmid encoding aerobactin. Infect Immun 54:603–608.

70 Llobet E, Tomas JM, Bengoechea JA. 2008. Capsule polysaccharide is a bacterial decoy for antimicrobial peptides. Microbiology 154:3877–3886.

